# Cholinergic interneurons mediate cocaine extinction through similar plasticity across medium spiny neuron subtypes

**DOI:** 10.1101/2021.08.29.458113

**Authors:** Weston Fleming, Junuk Lee, Brandy A. Briones, Scott Bolkan, Ilana B. Witten

**Affiliations:** Princeton Neuroscience Institute, Princeton University, Princeton, New Jersey, USA; Department of Psychology, Princeton University, Princeton, New Jersey, USA

## Abstract

Cholinergic interneurons (ChINs) in the nucleus accumbens (NAc) have been implicated in the acquisition and extinction of drug associations, as well as related plasticity in medium spiny neurons (MSNs). However, since most previous work has relied on artificial manipulations, if and how endogenous patterns of cholinergic signaling relate to drug associations is unknown. Moreover, despite great interest in the opposing effects of dopamine on MSN subtypes, whether ChIN-mediated effects are similar or different across MSN subtypes is also unknown. Here, we find that endogenous acetylcholine event frequency during extinction negatively correlates with the strength and persistence of cocaine-context associations across individuals, consistent with effects of artificial manipulation of ChIN activity during extinction. Moreover, ChIN activation during extinction produces a reduction in excitatory synaptic strength on both MSN subtypes, similar to the effect of multiple extinction sessions in the absence of ChIN manipulations. Together, our findings indicate that natural variation in NAc acetylcholine may contribute to individual differences in drug-context extinction by modulating glutamatergic presynaptic strength similarly at both D1R and D2R MSN subtypes.

**Graphical abstract:** 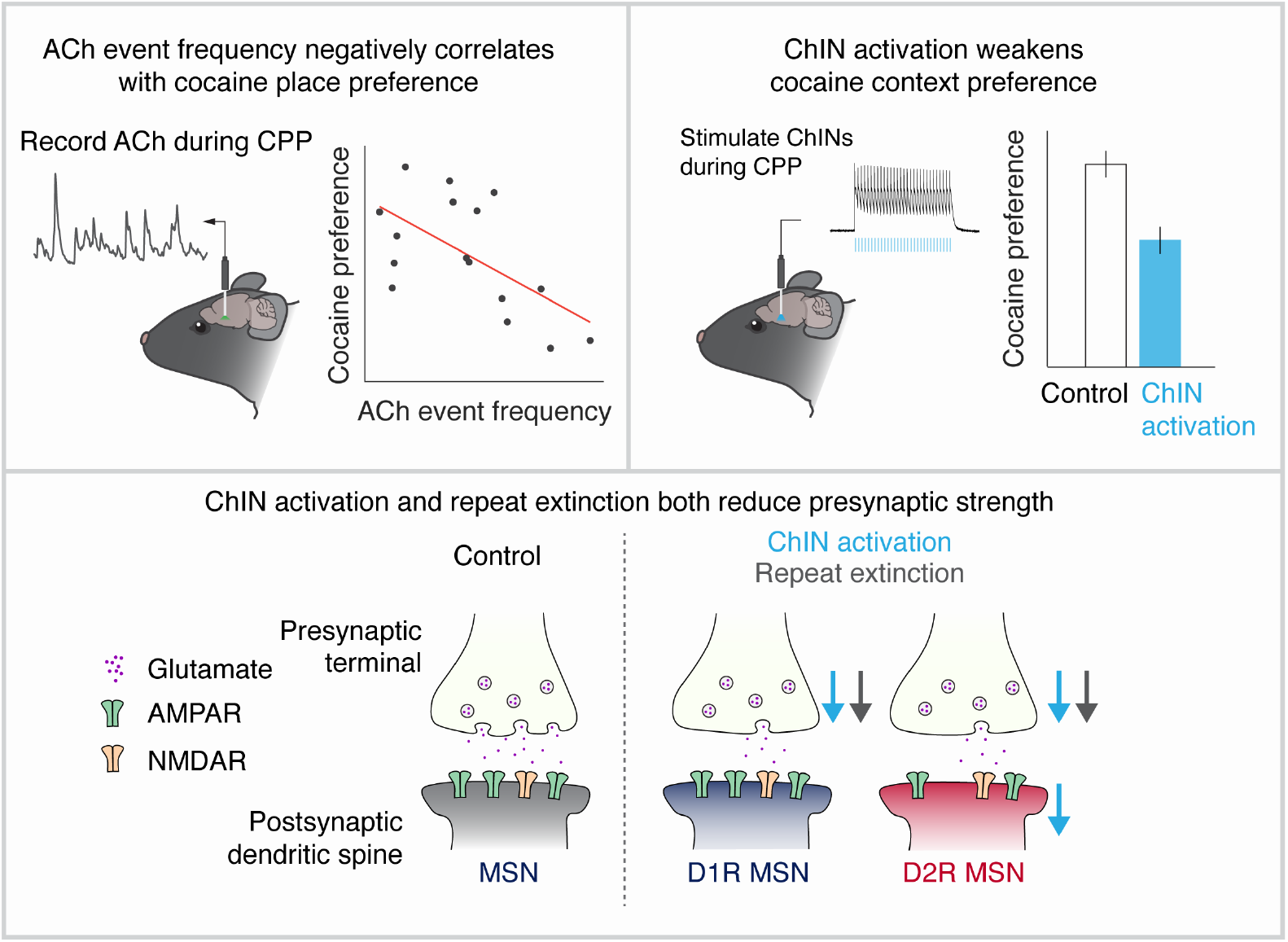

## Introduction

Animals must update reward-context associations when reward is no longer present, a process known as extinction. Understanding the neural mechanisms of extinction is not only of fundamental relevance, but also of potential clinical relevance. Drugs of abuse like cocaine can form persistent drug-context associations, and understanding the mechanisms that underlie their extinction may offer insight to treatments that promote abstinence.

Some of the neural processes that contribute to the formation and extinction of drug-context associations are believed to occur in the nucleus accumbens (NAc). Drugs of abuse generate high levels of dopamine release in the NAc. As a result, the role of dopamine in addiction-related NAc plasticity has been studied extensively (Britt et al., 2012; Kourrich et al., 2007; MacAskill et al., 2014; Pascoli et al., 2014; Robinson et al., 2001).

In addition, there is growing appreciation for a complementary role of acetylcholine (ACh) — a comparatively understudied neuromodulator in the NAc — in regulating plasticity and drug-context learning (Aitta-Aho et al., 2017; Collins et al., 2019; Lee et al., 2016; Witten et al., 2010). However, since previous work has relied primarily on artificial manipulation of cholinergic interneurons (ChINs) whether natural variation in ACh signaling across individuals is predictive of drug associations and their persistence is unknown.

Another open question is whether and how ChINs differentially modulate the output populations of the NAc to drive extinction of drug-context associations. The NAc is composed principally of medium spiny neurons (MSNs), which function as the structure’s primary output and can be classified into two subtypes by the dopamine receptor expressed (either D1R or D2R). Dopamine drives differential forms of plasticity in these two MSN subtypes (Gerfen et al., 1990; Iino et al., 2020; Lee et al., 2021; Shen et al., 2008), a phenomenon believed to contribute to the distinct and sometimes opposing roles these subpopulations have in reward-related behaviors (Bock et al., 2013; Cole et al., 2018; Durieux et al., 2009; Gallo et al., 2018; Hikida et al., 2010; Kravitz et al., 2012; Lobo et al., 2010; O’Neal et al., 2020).

However, whether *in vivo* ChIN activity also differentially affects plasticity at D1R vs D2R MSNs is unknown. It is possible that ChIN activity produces MSN subtype-specific plasticity, given the differential muscarinic receptor expression profiles across MSN cell types (Bernard et al., 1992; Lim et al., 2014; Yan et al., 2001), and given ACh’s ability to mediate dopamine release onto MSNs (Cachope et al., 2012; Threlfell et al., 2012; Yorgason et al., 2017), which is in turn thought to mediate opposing plasticity (Gerfen et al., 1990; Iino et al., 2020; Lee et al., 2021; Shen et al., 2008). Alternatively, ChINs may drive similar changes across MSN subtypes, given that MSNs are driven by input from glutamatergic projections (Britt et al., 2012; Phillipson and Griffiths, 1985) that themselves express muscarinic receptors (Hernández-Echeagaray et al., 1998; Higley et al., 2009; Sugita et al., 1991), and that are inhibited similarly by muscarinic activation *ex vivo* (Ding et al., 2010).

To address these gaps, we first used an ACh indicator to demonstrate that the frequency of ACh events is predictive of the strength and persistence of cocaine-context associations across individuals. This extends a previous observation (Lee et al., 2016) that artificial activation of ChINs enhances cocaine-context extinction by demonstrating a role for natural variations in ACh in mediating individual differences. In addition, we show that ChIN activation during a single extinction session reduces glutamatergic presynaptic strength at both MSN subtypes, while also reducing postsynaptic strength specifically at D2R MSNs. This effect is similar to plasticity observed from multiple days of extinction in the absence of a ChIN manipulation, implying that elevated ChIN activity hastens extinction by accelerating these naturally occurring plasticity mechanisms. Taken together, our data suggests that variations in ChIN activity may mediate individual differences in the extinction of cocaine-context associations by modulating the strength of glutamatergic inputs onto both D1R and D2R MSNs.

## Results

### Repeat extinction following cocaine CPP reduces excitatory transmission similarly at both D1R and D2R MSNs

Before characterizing the correlates and consequences of NAc ACh during cocaine-context extinction, we first sought to identify the extinction-dependent changes in glutamatergic synaptic strength at D1R and D2R MSNs.

To study plasticity changes related to drug-context extinction, we conditioned and extinguished mice through a cocaine conditioned place preference (CPP). During the first day of the CPP (“Baseline”), mice had access to both zones of the CPP chamber. The next 2 days, mice underwent conditioning, during which one zone was paired with cocaine (15 mg/kg, i.p.) and the other with saline. One group of mice (“Repeat extinction group”) underwent 4 subsequent days of extinction testing (“Tests 1-4”) to assess and extinguish their preference. To control for plasticity effects that may occur over time independent of extinction experience, a separate group (“No extinction”) underwent the conditioning paradigm, but did not undergo additional testing after conditioning (**Figure 1a**). As expected, mice that underwent extinction showed a preference for the cocaine zone on Test 1, which decreased on Tests 2-4 (**Figure 1b**). Cocaine-mediated locomotor activation during cocaine conditioning did not differ between repeat extinction and no extinction groups (**Supplementary figure 1**).

**Figure 1.**
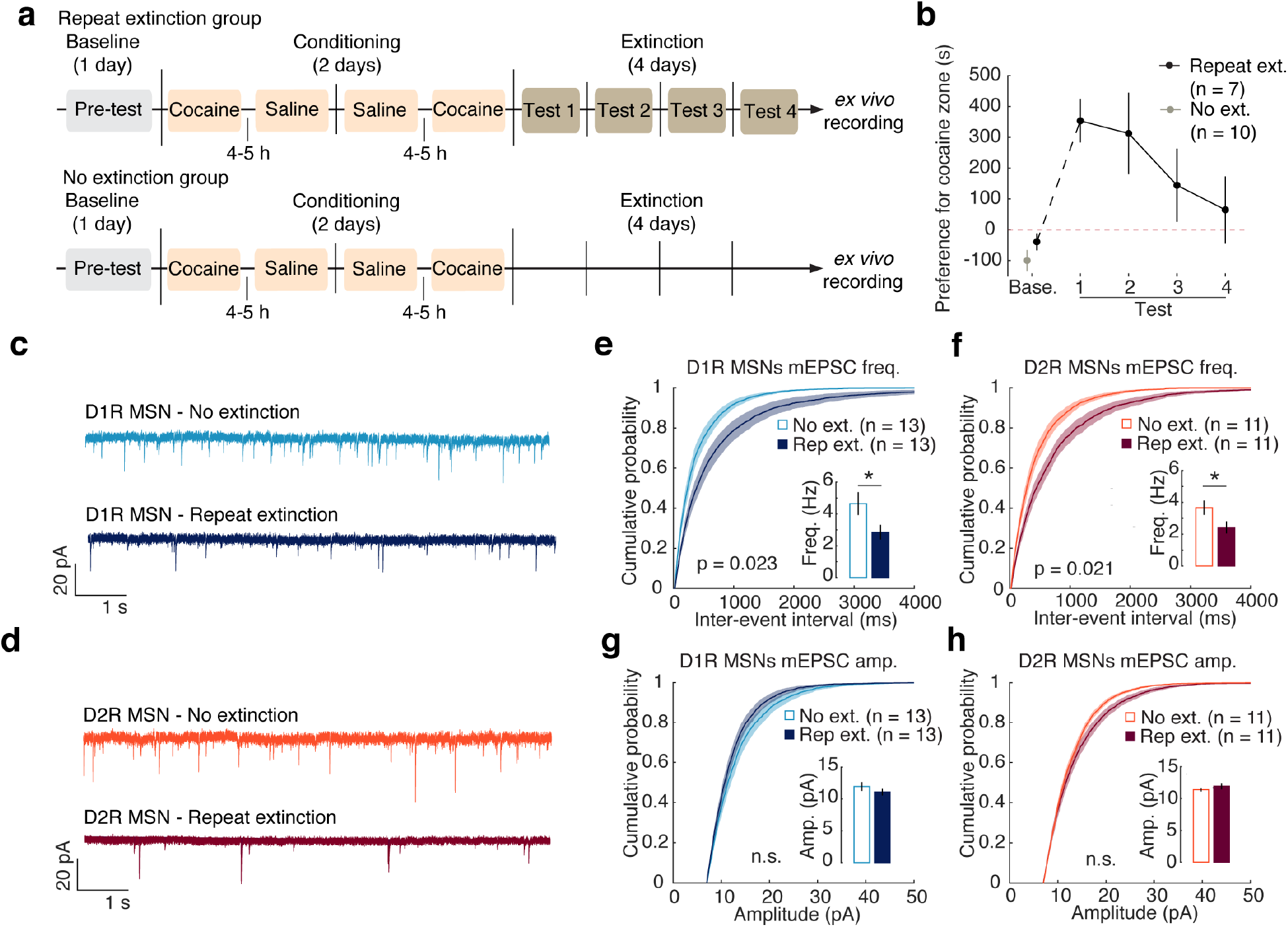
Repeat extinction following cocaine CPP reduces excitatory transmission at both NAc MSN subtypes. **a.** Timeline of CPP experiments followed by either repeat extinction testing or no extinction. For the repeat extinction group, brain slices were collected on Day 7 immediately after Test 4. For the no extinction group, brain slices were collected on Day 7 at approximately the same time of day as brain slice collection for the repeat extinction group. **b.** Preference for the cocaine zone, measured as the difference in time spent on the cocaine zone minus time spent on the saline zone. Mice from the repeat extinction group preferred the cocaine-paired chamber more on Test 1 than on Baseline, and this preference extinguished across days (F_(4,24)_ = 6.388, p = 0.001 for session in one-way repeated measures ANOVA with the 5 sessions as fixed effect and mouse as random effect.). **c.** Example traces of whole-cell, voltage-clamp mEPSCs from D1R MSNs from mice from repeat extinction and no extinction groups. Scale bars are 20 pA and 1 s. **d**. Same as **c**, but example traces from D2R MSNs. **e.** Cumulative probability of interevent intervals for mEPSCs at D1R MSNs. Repeat extinction decreased mEPSC frequency in D1R MSNs compared to mice that did not undergo extinction testing (Effect size d = 0.989, p = 0.023 for group in a linear, mixed-effects regression on frequency with group as a fixed effect and cell as random effect. Inset: median frequency of mEPSCs. Repeat extinction group, n = 13 cells, 4 mice, 2.852 Hz ± 0.460. No extinction group, n = 13 cells, 4 mice, 4.658 Hz ± 0.719). **f.** Cumulative probability of interevent intervals for mEPSCs at D2R MSNs. Repeat extinction decreased mEPSC frequency in D2R MSNs compared to mice that did not undergo extinction testing (Effect size d = 1.118, p = 0.021 for group in a linear, mixed-effects regression on frequency with group as a fixed effect and cell as random effect. Inset: median frequency of mEPSCs. Repeat extinction group, n = 11 cells, 4 mice, 2.430 Hz ± 0.379. No extinction group, n = 11 cells, 4 mice, 3.667 Hz ± 0.451). **g.** Cumulative probability of amplitudes for mEPSCs at D1R MSNs. Repeat extinction testing did not affect mEPSC amplitude in D1R MSNs compared to mice that did not undergo extinction testing (Effect size d = 0.429, p = 0.304 for group in a linear, mixed-effects regression on amplitude with group as a fixed effect and cell as random effect. Inset: median amplitude of mEPSCs. Repeat extinction group, n = 13 cells, 4 mice, 11.086 pA ± 0.541. No extinction group, n = 13 cells, 4 mice, 11.949 pA ± 0.686). **h.** Cumulative probability of amplitudes for mEPSCs at D2R MSNs. Repeat extinction testing did not affect mEPSC amplitude in D2R MSNs compared to mice that did not undergo extinction testing (Effect size d = -0.432, p = 0.345 for group in a linear, mixed-effects regression on amplitude with group as a fixed effect and cell as random effect. Inset: median amplitude of mEPSCs. Repeat extinction group, n = 11 cells, 4 mice, 11.924 pA ± 0.4836. No extinction group, n = 11 cells, 4 mice, 11.375 pA ± 0.289).

To assess how extinction affected glutamatergic synaptic strength at MSNs in NAc medial shell, brain slices were taken from both groups on Day 7 and mini excitatory postsynaptic currents (mEPSCs) were recorded. We used drd1a-tdTomato and drd2-GFP mice to target recordings to each MSN subtype after first validating that the lines display high degrees of specificity and penetrance (Supplementary figure 2).

Repeat extinction significantly reduced mEPSC frequencies at both D1R and D2R MSNs compared to mice that did not undergo extinction (**Figure 1c-f**; Effect size d = 0.989, p = 0.023 for repeat extinction vs no extinction group in a linear, mixed-effects regression, LMER, on D1R frequency; Effect size d = 1.118, p = 0.021 for group in an LMER on D2R frequency). In contrast, there was no significant effect on amplitudes at either population (**Figure 1g-h**; p = 0.304 for repeat extinction vs no extinction group in an LMER on D1R amplitudes; p = 0.345 for group in an LMER on D2R amplitudes). These findings suggest that extinction experience drives a reduction of presynaptic strength in excitatory transmission onto both NAc D1R and D2R MSNs.

### Higher ACh event rate during cocaine extinction is associated with lower cocaine preference and persistence

Before characterizing how ChIN activity contributes to the observed plasticity in each MSN cell-type during extinction, we examined how endogenous ACh release relates to behavior. Towards this end, we injected a GRABACh sensor (AAV2/9-hSyn-GACh3.0) (Jing et al., 2020) into the medial shell of NAc in order to record ACh levels during a cocaine CPP (**Figure 2a-b**), after first confirming *ex vivo* that the indicator was selectively activated by ACh (**Figure 2c**). Mice underwent a cocaine CPP followed by 4 days of extinction (“Tests 1-4”; **Figure 2d**). During cocaine conditioning, there was a large decrease in ACh event frequency and amplitude compared to saline conditioning (**Figure 2e**; **Supplementary figures 3-4**). Following conditioning, compared to baseline, mice showed a significant preference for the cocaine zone that decreased with subsequent tests, indicating extinction learning (**Supplementary figure 5**).

**Figure 2.**
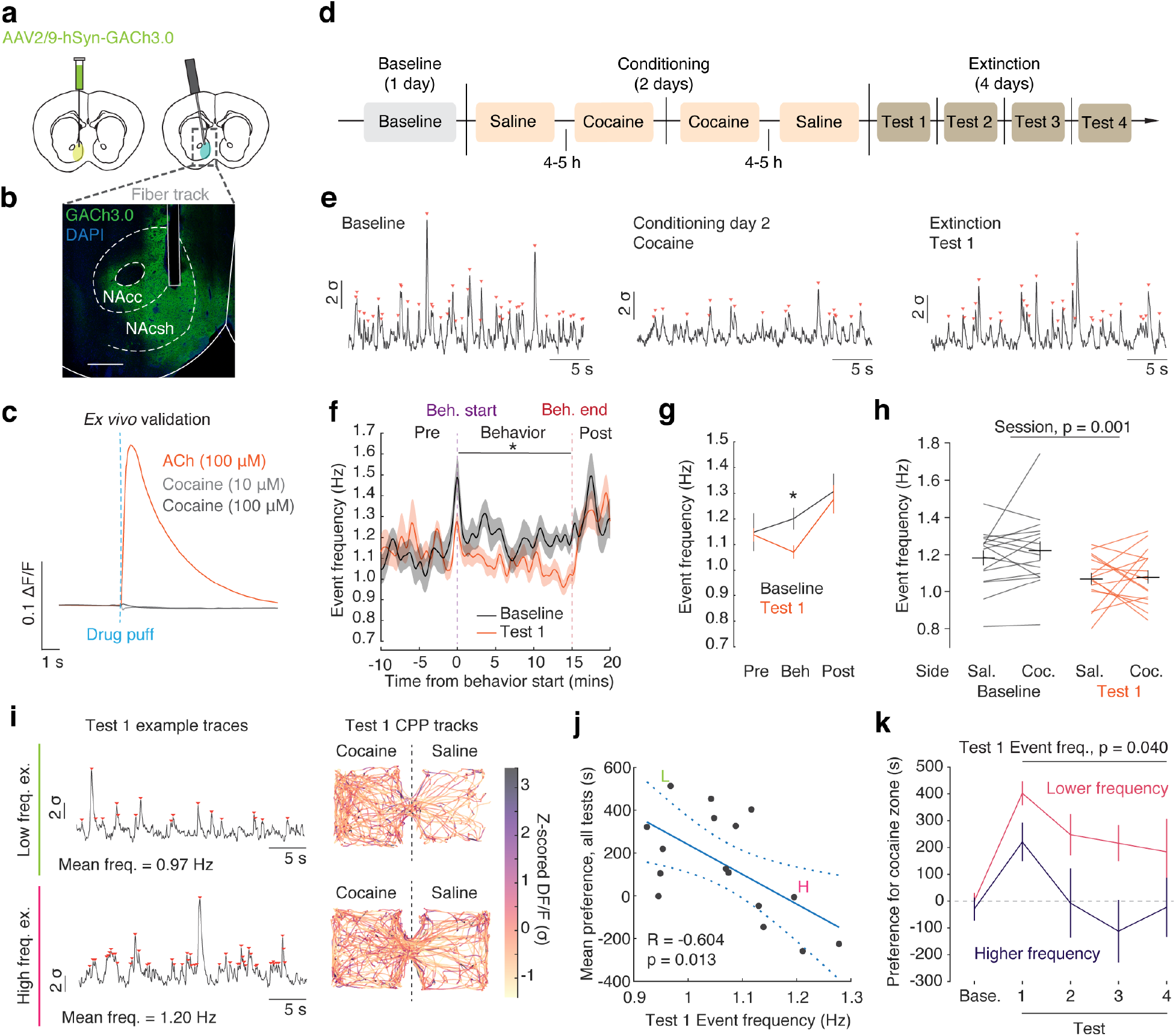
Mice with higher ACh event rate during cocaine extinction exhibit less cocaine preference. **a.** AAV2/9- hSyn-GACh3.0 virus was injected into the NAc of mice. Fibers were implanted above the injection sites for unilateral recordings. **b.** Representative image showing GACh3.0 expression in NAc with fiber placement. **c.** *Ex vivo* imaging shows tissue expressing GACh3.0 is responsive to acetylcholine (ACh) puff but not cocaine. **d.** Timeline of CPP experiments. **e.** Example z-scored dF/F of GACh3.0 fiber photometry traces from the same mouse with demarcated detected events (red triangles) during Baseline, Cocaine conditioning 2, and Test 1. **f.** Mean event frequency calculated with a 30 s moving window before, during, and after Baseline (grey) and Test 1 (orange) sessions. **g.** Mean event frequencies were significantly decreased during behavior testing (Beh), but not in periods before (Pre) or after (Post) behavior (p = 0.893 for pre-behavior period pairwise t-test of Baseline and Test 1; p = 0.013 for behavior period pairwise t-test of Baseline and Test 1; p = 0.730 for post-behavior period pairwise t-test of Baseline and Test 1). **h.** Mean event frequency is not different between cocaine- and saline-conditioned zones on Baseline or Test 1 (F_(1,45)_ = 12.683, p = 0.001 for session (Test 1 vs Baseline); F_(1,45)_ = 0.468, p = 0.498 for chamber side (saline vs cocaine); F_(1,45)_ = 0.168, p = 0.684 for session*chamber side in repeated measures ANOVA on mean event frequency with session, chamber side, and their interaction as fixed effects and mouse as random effect). **i.** Left: z-scored dF/F of GACh3.0, recorded during Test 1 from two example mice with either low event frequency (top) or high event frequency (bottom). Right: Example CPP test tracks from the same mice during Test 1, colored with z-score value of ACh3.0 fluorescence. Scale bar is constrained to 1st and 99th percentile values of z-score values from the two mice. **j.** Across individuals, event frequency on Test 1 is significantly negatively correlated with mean preference for the cocaine zone on Tests 1-4 (R = -0.604; p = 0.013). **k.** Mean preference for the cocaine zone when mice are median split by Test 1 frequency (Lower frequency, pink, n = 8; Higher frequency, purple, n = 8). Event frequency on Test 1 is significantly predictive of preference on Tests 1-4 (F_(1,14)_ = 5.10, p = 0.040 for median split group in repeated measures ANOVA on preference with group, test number, and their interaction as fixed effects and mouse as random effect).

On the first extinction test after conditioning (Test 1), there was a significant decrease in ACh event frequency compared to baseline (Effect size d = 0.725, p = 0.011 for pairwise t-test of Baseline and Test 1; **Figure 2f-g**; **Supplementary figure 5**). This decrease was only evident in the behavioral chamber and not during the periods immediately before or after, when the mouse was in its homecage (p = 0.893 for 10 min pre-behavior period pairwise t-test of Baseline and Test 1; p = 0.730 for 5 min post-behavior period pairwise t-test of Baseline and Test 1; **Figure 2f-g**). Event frequency was similar regardless of chamber side (**Figure 2h**; **Supplementary figures 6**), and event frequency showed similar modulation during entries to the cocaine and saline zones (**Supplementary figure 7**).

Given that artificial manipulation of ChINs is thought to modulate cocaine CPP extinction (Lee et al., 2016), and given that we observed considerable variation in ACh event rates across individuals (**Figure 2h-i**), we wondered if individual differences in naturally occuring ACh may be predictive of the strength and persistence of cocaine-context associations. Indeed, we found that ACh event frequency on Test 1 was negatively correlated with cocaine preference across days (correlation coefficient of Test 1 frequency vs mean preference over Tests 1-4, R = -0.604; p = 0.013; F(1,14) = 5.10, p = 0.040 for for median split group in repeated measures ANOVA on preference; **Figure 2j-k**; **Supplementary figure 5**). Thus, heightened ACh at the first extinction test is associated with weaker and less persistent cocaine preference.

In addition to this post-conditioning decrease in ACh event frequency, we also observed a post-conditioning change in the relationship between an animal’s speed and the timing of ACh events. Specifically, while speed was similar before versus after conditioning, especially on the cocaine side (**Figure 3a**), and ACh events had little relation to speed before conditioning (**Figure 3b**), there was a pronounced decrease in speed at the time of ACh events after versus before conditioning (**Figure 3c-d**; p = 7.4*10^-5^ for Baseline vs Test 1 session in repeated measures ANOVA on change in speed around event peak). This relationship was similar in the cocaine and saline zone (**Figure 3c**), but was not evident in mice that received saline-only instead of cocaine conditioning (**Supplementary figure 8**). Similarly, after conditioning, higher speeds were more negatively associated with ACh event frequencies (**Figure 3e**). Interestingly, mice with a more negative relationship between speed and ACh event rates after conditioning exhibited less persistent cocaine preference (**Figure 3e-f**, **Supplementary figure 9**; F(1,14) = 8.337, p = 0.012 for median split group in repeated measures ANOVA). Mean speed on Test 1 did not correlate with mean ACh event frequency or cocaine zone preferences on Tests 1-4 (**Supplementary figure 10**), indicating that overall speed is not predictive of preference or ACh events.

**Figure 3.**
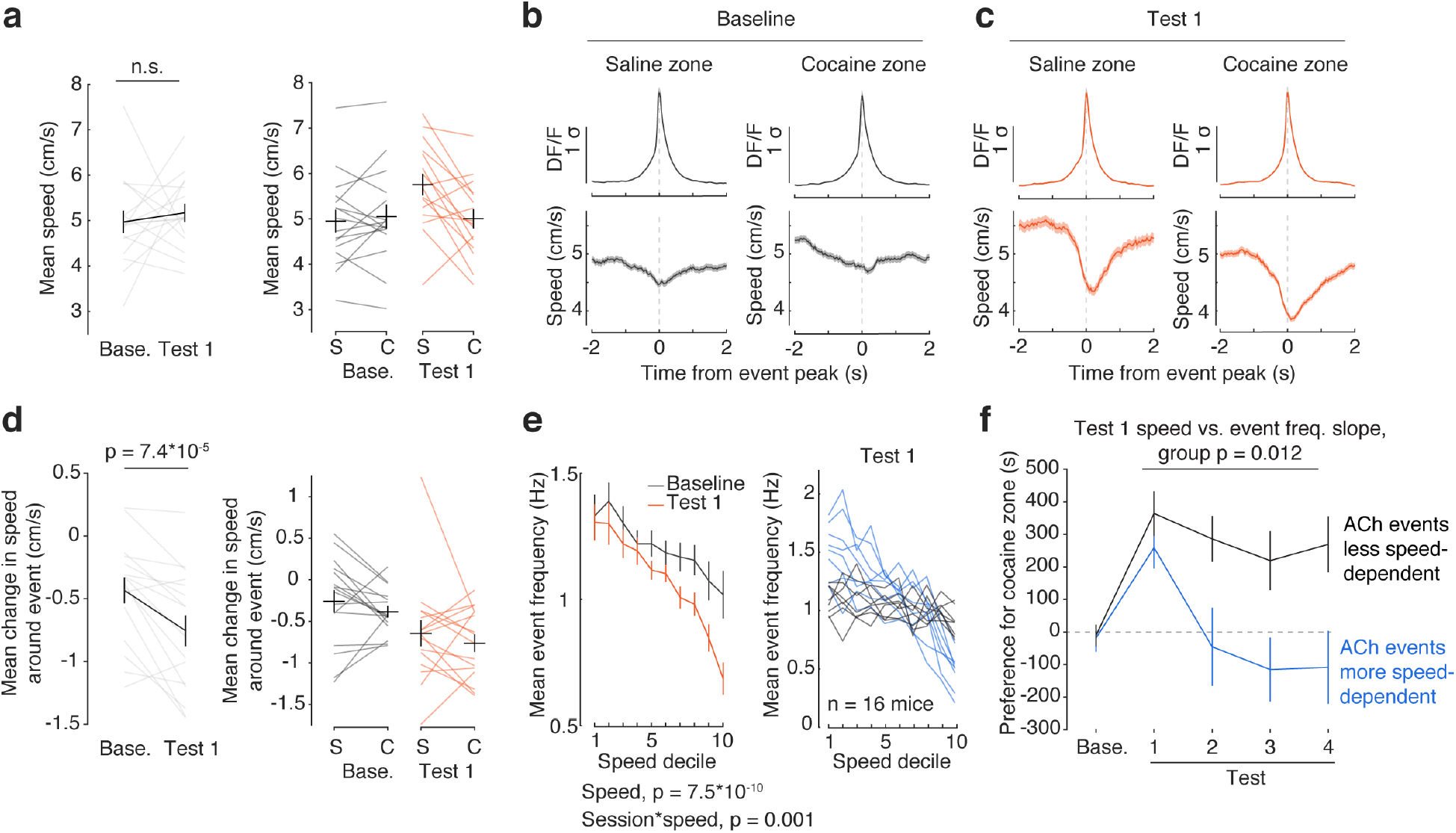
Mice with less persistent cocaine preference have a negative relationship between ACh events and speed after conditioning. **a.** Mean speed during Baseline and Test 1 sessions (left) and broken down by chamber side (right). Mice maintain similar speeds between Baseline and Test 1 and regardless of chamber side (F_(1,45)_ = 2.875, p = 0.097 for session (Test 1 vs Baseline); F_(1,45)_ = 2.185, p = 0.146 for chamber side (saline vs cocaine); F_(1,45)_ = 3.741, p = 0.059 for session*chamber side in repeated measures ANOVA on mean speed with session, chamber side, and their interaction as fixed effects and mouse as random effect). **b.** Top: Mean z-scored dF/F trace of GACh3.0 signal centered at peak time for all significant events across all mice in the saline zone (left) and cocaine zone (right) during the Baseline test. Bottom: Mean speed across all events from all mice over the same time period in the saline zone (left) or cocaine zone (right) during Baseline (n = 8589 events in saline zone, 8610 events in cocaine zone). **c.** Same as **b**, but for the first extinction test following cocaine CPP (Test 1; n = 5057 events in saline zone, 10286 events in cocaine zone). **d.** Mean change in speed around ACh events during Baseline and Test 1 (left) and broken down by chamber side (right). Events coincide with larger decreases in speed on Test 1 than baseline (F_(1,45)_ = 19.035, p = 7.41*10^-5^ for session in repeated measures ANOVA on change in speed with session (Baseline vs Test 1), chamber side (saline vs cocaine), and their interaction as fixed effects and mouse as random effect). Changes in speed around time of events were calculated as mean speed in a 1.5 s window beginning 0.5 s before event peak and ending 1 s after event peak, baseline subtracted with the mean speed in a 1 s window starting 2 s before the event peak. **e.** Left: Mean event frequency by speed decile during Baseline and Test 1. Speed deciles are calculated for each mouse across all speed data from Baseline and Test 1 sessions. Event frequency is negatively modulated by speed, and to a greater extent at Test 1 than at Baseline (Effect size d = -0.753, p = 7.5*10^-10^ for speed; effect size d = 0.410, p = 0.001 for session*speed in an LMER with session (Baseline vs Test 1), speed decile, and their interaction as fixed effects and mouse as random effect). Right: Mean event frequency by speed decile on Test 1 for individual mice. Individual traces are colored by median split of slope to highlight differences in speed dependency (black: less negative speed vs. frequency slope; blue: more negative speed vs. frequency slope). **f.** Mean preference for the cocaine zone when mice are median split by the slopes of the speed decile vs. Test 1 mean event frequency (Less speed dependence, black, n = 8; More speed dependence, blue, n = 8). Speed decile slope is significantly predictive of preference on Tests 1-4 (F_(1,14)_ = 8.337, p = 0.012 for median split group in repeated measures ANOVA with group, test number, and their interaction as fixed effects and mouse as random effect).

Taken together, this means that stronger cocaine preference across individuals is predicted by low ACh event rates after conditioning (**Figure 2j-k**), and a tendency for the remaining events to occur at times of higher speed (**Figure 3e-f**).

### Activation of ChINs during extinction generates reductions in excitatory synaptic transmission across MSN subtypes, mimicking changes that occur as a consequence of repeated extinction

Given that high ACh event rates were predictive of weaker cocaine CPP strength and persistence (**Figure 2j-k**), and that extinction is associated with synaptic plasticity on both D1R and D2R neurons (**Figure 1e-f**), we next sought to determine if activation of ChINs during a single extinction session would decrease cocaine preference and reproduce the observed plasticity across MSN subtypes that was caused by multiple days of extinction.

To virally target ChINs for manipulation during behavior, and to identify MSNs by subtype during subsequent *ex vivo* electrophysiology experiments, we used ChAT::IRES-Cre mice crossed with either drd1a-tdTomato or drd2-GFP mice. An AAV2/5 virus expressing Cre-dependent ChR2-YFP was injected into the medial shell of NAc, and optical fibers were implanted bilaterally above the structure (**Figure 4a-b**; **Supplementary figure 11**). Mice underwent the same cocaine CPP as described in the fiber photometry experiment, and on day 4 (“Test”) underwent extinction testing (**Figure 4c**). For the duration of this test, one group of mice received phasic optogenetic activation of ChINs bilaterally as they explored the chamber (“ChIN activation group”). As we did not observe differences in ACh event rate based on chamber side during the extinction test (**Figure 2h**), we applied spatially uniform activation of ChINs (15 Hz, 2 s on, 2 s off; **Figure 4b**).

**Figure 4.**
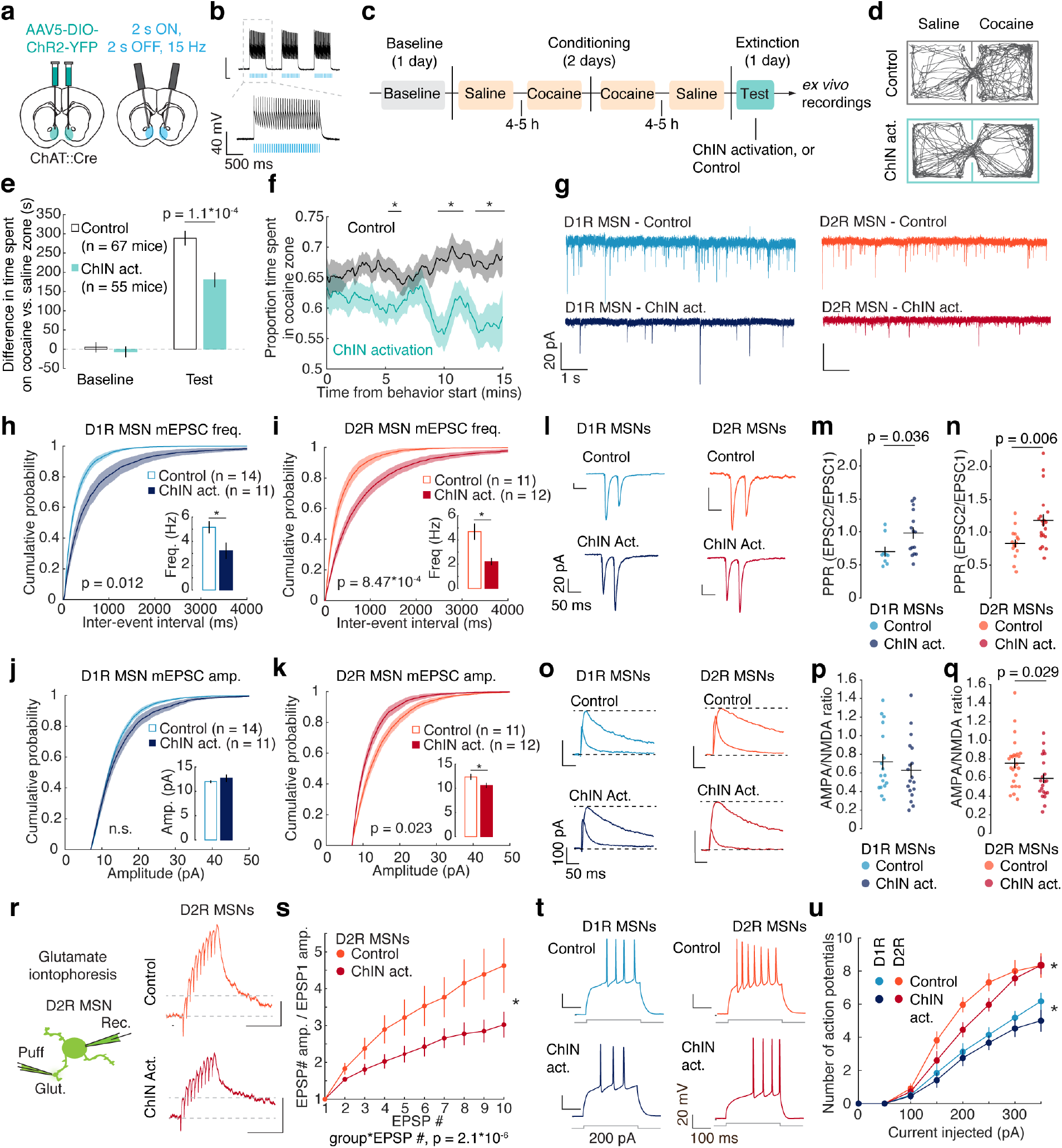
ChIN activation during cocaine extinction weakens presynaptic glutamatergic synapses in both MSN subtypes, with D2R MSN-specific postsynaptic effects. **a.** Cre-dependent AAV5-ChR2-YFP virus was injected bilaterally into the NAc of ChAT::IRES-Cre / drd1a-tdTomato mice or ChAT::IRES-Cre / drd2-GFP mice. Fibers were implanted above the injection sites. **b.** Whole-cell current clamp recordings to confirm the functionality of ChR2 in NAc ChINs. **c.** Timeline of CPP experiment. Both groups underwent a CPP, and ChIN activation group received optogenetic stimulation during extinction testing (447 nm, 15 Hz, 5 ms pulse duration, 2 s light on interleaved with 2 s light off). Immediately after the test, brain slices were collected for *ex vivo* recordings. **d.** Example locomotor tracks during extinction testing for a mouse that received ChIN activation and a control mouse. **e.** Activation of ChINs during extinction testing reduced preference for the cocaine-paired chamber compared to control mice (F_(1,118)_ = 273.282, p = 2*10^-16^ for test condition. F_(1,118)_ = 10.884, p = 0.001 for test condition*light condition interaction; F_(1,118)_ = 10.043, p = 0.002 for light condition; F_(1,118)_ = 0.307, p = 0.581 for genotype; F_(1,118)_ = 0.689, p = 0.408 for genotype*test condition; Multi- factor, repeated measures ANOVA with transgenic mouse line, group (ChIN activation vs control), session (Baseline vs Test), and their interactions as fixed effects, and mouse as random effect; p = 0.550 for post-hoc test of group (ChIN activation vs control) on Baseline, p = 1.1*10^-4^ for group on Test, Holm-Sidak’s post hoc test). **f.** Mean proportions of time spent in the cocaine zone in the Test session for Control and ChIN activation groups, plotted as moving means with a centered 2 min window. Activation of ChINs during extinction testing reduced preference as a function of time. (p = 0.025 for group*time interaction in an LMER with group (ChIN activation vs control), time (1 min non-overlapping, unsmoothed bins), and their interaction as fixed effects and mouse as random effect; p < 0.05 at minutes 6, 10-11, 13- 15 in post-hoc pairwise t-tests of each minute between groups with Holm’s correction). **g.** Example traces of whole-cell, voltage-clamp mEPSCs from D1R and D2R MSNs from mice that received ChIN activation during extinction testing and control mice. **h.** Cumulative probability of inter-event intervals for mEPSCs at D1R MSNs. ChIN activation decreased mEPSC frequency at D1R MSNs compared to controls (Effect size d = 1.139, p = 0.012 for group in an LMER on frequency with group (ChIN activation vs control) as a fixed effect and cell as random effect. Inset: median frequency of mEPSCs. Control group, n = 14 cells, 13 mice, 5.16 ± 0.50 Hz. ChIN activation group, n = 11 cells, 10 mice, 3.23 ± 0.66 Hz). **i.** Cumulative probability of interevent intervals for mEPSCs at D2R MSNs. ChIN activation decreased mEPSC frequency at D2R MSNs compared to controls (Effect size d = 1.697, p = 8.47*10^-4^ for group in an LMER on frequency with group (ChIN activation vs control) as a fixed effect and cell as random effect. Inset: median frequency of mEPSCs. Control group, n = 11 cells, 6 mice, 4.69 ± 0.67 Hz. ChIN activation group, n = 11 cells, 6 mice, 2.22 ± 0.29 Hz). **j.** Cumulative probability of amplitudes for mEPSCs at D1R MSNs. ChIN activation did not affect mEPSC amplitude in D1R MSNs compared to controls (Effect size d = 0.336, p = 0.429 for group in an LMER on amplitude with group (ChIN activation vs control) as a fixed effect and cell as random effect. Inset: median amplitude of mEPSCs. Control group, n = 14 cells, 13 mice, 11.98 ± 0.28 pA. ChIN activation group, n = 11 cells, 10 mice, 12.69 ± 0.79 pA). **k.** Cumulative probability of amplitudes for mEPSCs at D2R MSNs. ChIN activation reduced mEPSC amplitudes in D2R MSNs compared to controls (Effect size d = -1.067, p = 0.023 for group in an LMER on amplitude with group (ChIN activation vs control) as a fixed effect and cell as random effect. Inset: median amplitude of mEPSCs. Control group, n = 11 cells, 9 mice, 12.35 ± 0.64 pA. ChIN activation group, n = 11 cells, 6 mice, 10.58 ± 0.55 pA). **l.** Representative EPSCs in D1R and D2R MSNs in response to interpulse intervals of 50 ms in control mice and mice that received ChIN activation during extinction testing. Scale bars: 50 ms and 20 pA. **m.** Compared to controls, ChIN activation increased paired-pulse ratio at D1R MSNs (Effect size d = 0.994, p = 0.036, two-tailed t test; Control group, n = 9 cells, 0.700 ± 0.0727. ChIN activation group, n = 15 cells, 0.980 ± 0.086). **n.** Compared to controls, ChIN activation increased paired-pulse ratio at D2R MSNs (Effect size d = 1.037, p = 0.006, two-tailed t test; Control group, n = 15, 0.828 ± 0.059. ChIN activation group, n = 20, 1.175 ± 0.093). **o.** Representative traces of AMPA and NMDA receptor- mediated currents measured in voltage clamp at +40 mV. Scale bars: 50 ms and 100 pA. **p.** ChIN activation did not affect AMPA/NMDA ratio at D1R MSNs compared to controls (Effect size d = 0.270, p = 0.431, two-tailed t test; Control group, n = 17 cells, 0.719 ± 0.083. ChIN activation group, n = 18 cells, 0.629 ± 0.078). **q.** ChIN activation reduced AMPA/NMDA ratio at D2R MSNs compared to controls (Effect size d = 0.671, p = 0.029, two-tailed t test; Control group, n = 25 cells, 0.756 ± 0.052. ChIN activation group, n = 21 cells, 0.591 ± 0.050). **r.** Schematic of iontophoretic application of glutamate to MSN dendrites and example traces of postsynaptic response to iontophoretic glutamate application at dendrites of D2R MSNs of a control mouse (top) and a mouse that received ChIN activation (bottom) during extinction test. **s.** Group mean magnitudes of EPSP response to each application of glutamate, normalized to the first EPSP. Cells from mice that had received ChIN activation during extinction show significantly less summation of EPSPs (Effect size d = 0.804, p = 2.1*10^-6^ for group*EPSP number in an LMER with group (ChIN activation vs control), EPSP number, and their interaction as fixed effects and cell as random effect; control n = 10 cells; ChIN activation n = 7 cells). **t.** Representative traces of evoked action potentials during a 300 ms, 200 pA current injection in D1R and D2R MSNs from mice that received ChIN activation (ChIN act.), and controls that received no light, during extinction testing. Scale bars: 300 ms and 20 mV. **u.** Plots of mean number of evoked action potentials vs. injected current in D1R MSNs (blue) and D2R MSNs (red). For both MSN subtypes, there was a significant interaction effect between group and current (Effect size d = 0.281, p = 0.046 for current *group interaction for D1R MSNs in an LMER with group (ChIN activation vs control), injected current, and their interaction as fixed effects and cell as random effect; p = 0.044 in D1R MSNs for pairwise group comparison on slope of current-spike response; Effect size d = 0.322, p = 0.040 for current*group interaction for D2R MSNs in an LMER with group (ChIN activation vs control), injected current, and their interaction as fixed effects and cell as random effect; p = 0.038 in D2R MSNs for pairwise group comparison on slope of current-spike response). All error bars are SEM.

Consistent with our indicator data (**Figure 2**), as well as previous optogenetic manipulations (Lee et al., 2016), mice that received optogenetic activation of ChINs during testing showed significantly less preference for the cocaine zone compared to controls (**Figures 4d-e**) (F(1,118) = 10.884, p = 0.001 for session*group interaction in multi-factor, repeated measures ANOVA on preference; p = 1.1*10^-4^ for group (ChIN activation vs control) on Test session, pairwise t-test with Holm-Sidak’s correction). The difference in preference between the ChIN activation and control groups emerged during the course of the Test session, suggesting that ChIN activation accelerates extinction (**Figure 4f**; p = 0.025 for group*time interaction in an LMER; p < 0.05 at minutes 6, 10-11, 13-15 in pairwise t-test between groups (ChIN activation vs control) with Holm’s correction). There was no effect of mouse line on cocaine preference or the effect of ChIN stimulation, and there was no difference in mean speed during the Test session between experimental groups (**Supplementary figures 12-13**).

Immediately following extinction testing, brain slices were collected for plasticity measurements (**Figure 4c**). We first measured mEPSCs to determine whether optogenetic activation of ChINs generated reductions in mEPSC frequency as seen following repeat extinction (**Figure 4g**). Indeed, compared to controls, mice who received ChIN activation during extinction testing had significantly lower mEPSC frequencies at both D1R and D2R MSNs (**Figure 4h-i**; **Supplementary figure 14**) (Effect size d = 1.139, p = 0.012 for group in an LMER on frequency for D1R MSNs; Effect size d = 1.697, p = 8.47*10^-4^ for group in an LMER on frequency for D2R MSNs). In addition, mice who received ChIN activation during extinction testing showed reduced mEPSC amplitudes at D2R MSNs (Effect size d = -1.067, p = 0.023 for group in an LMER on amplitude) but not at D1R MSNs (Effect size d = 0.336, p = 0.429 for group in an LMER on amplitude; **Figures 4j-k**). This suggests ChIN activation during extinction causes presynaptic weakening on both cell types and postsynaptic weakening on D2R MSNs.

To confirm the pre- and postsynaptic sites of these plasticity effects, we performed additional experiments in which glutamatergic afferents onto NAc MSNs were electrically stimulated and evoked EPSCs were measured. We first performed paired pulse ratio (PPR) measurements as an assay of presynaptic strength (Choi and Lovinger, 1997; Tang et al., 2001) (**Figure 4l**). Mice that received ChIN activation during extinction testing showed significant enhancement of PPRs at both D1R and D2R MSNs compared to controls (**Figure 4m-n**. Effect size d = 0.994, p = 0.036 for D1R MSNs, two-tailed t test; Effect size d = 1.037, p = 0.006 for D2R MSNs, two-tailed t test), consistent with the observed reduction in mEPSC frequencies (**Figure 4h-i**). Next, AMPA and NMDA receptor currents were measured at MSNs following extinction testing (**Figure 4o**). ChIN activation during extinction produced a significant reduction in AMPA/NMDA ratio compared to controls at D2R MSNs (Effect size d = 0.671, p = 0.029 for D2R MSNs, two-tailed t test) but not D1R MSNs (Effect size d = 0.270, p = 0.431 for D1R MSNs, two-tailed t test; **Figure 4p-q**), consistent with our observation that ChIN activation reduced mEPSC amplitudes only at D2R MSNs (**Figure 4k**). Taken together, these experiments suggest that ChIN activity promotes extinction of cocaine CPP through a combination of pre- and post-synaptic plasticity effects that dampen glutamatergic transmission at both D1R and D2R MSNs.

As another readout of postsynaptic plasticity, we also examined whether *in vivo* ChIN activation during extinction affected temporal summation of glutamatergic inputs. This experiment was motivated by a previous study which found that *ex vivo* muscarine treatment enhanced temporal summation in D2R MSNs (Shen et al., 2007) To test for such effects of ChIN activity *in vivo* during extinction, we repeatedly and rapidly iontophoretically delivered glutamate onto visually identified dendrites of D2R MSNs (**Figure 4r**). In contrast to the previous study, which performed the manipulation *ex vivo* (Shen et al., 2007), ChIN activation during extinction testing decreased temporal summation of EPSPs in D2R MSNs (Effect size d = 0.804, p = 2.1*10^-6^ for group*EPSP number in an LMER; **Figure 4s**). This further supports our observation that ChIN activation during extinction reduces the excitatory influence of glutamatergic signals arriving at D2R MSNs.

### Activation of ChINs during extinction produces minimal effects on membrane excitability in MSNs

We also examined if ChIN activity during extinction may affect MSN membrane excitability, in addition to the effects on synaptic strength (**Figure 4t**). Cells were recorded in current clamp at resting potentials of approximately -80 mV while current steps were injected (-200 pA to 350 pA, 50 pA intervals). In comparing the number of spikes evoked by the amount of current injected, there was a small but significant interaction between ChIN activation and current at both D1R and D2R MSNs, indicating a subtle decrease in excitability (**Figures 4u**) (Effect size d = 0.281, p = 0.046 for current*group interaction for D1R MSNs in an LMER; p = 0.044 in D1R MSNs for pairwise group comparison on slope of current-spike response; Effect size d = 0.322, p = 0.040 for current*group interaction for D2R MSNs in an LMER; p = 0.038 in D2R MSNs for pairwise group comparison on slope of current-spike response). There was no effect of ChIN activation on the action potential threshold, spike amplitude, first inter-spike interval, or afterhyperpolarization statistics for either MSN subtype (**Supplementary figure 15**). Moreover, ChIN activation decreased latency to spike in D1R MSNs (**Supplementary figure 15**), a form of increased excitability. Thus, while ChIN activation has profound effects in weakening glutamatergic synaptic strength, it produces only subtle effects on MSN membrane excitability.

## Discussion

Here, we first demonstrate that cocaine-context extinction decreases excitatory synaptic strength across MSN subtypes. During the initial extinction test, *in vivo* ACh event frequency is lower than before conditioning, and ACh event frequency is inversely related to the strength and persistence of cocaine-context associations. Consistent with these correlates, optogenetic activation of ChINs during extinction enhances extinction, while reducing glutamatergic synaptic strength at both D1R and D2R MSNs. Thus, these findings implicate an ACh-mediated reduction in glutamatergic synaptic strength across MSN subtypes as a key mechanism underlying cocaine-context extinction.

### Individual variability in ACh signaling predicts strength and persistence of cocaine-context associations

While there is growing appreciation that ACh plays an important role in drug learning, whether differences in the *in vivo* pattern of ACh across individuals are predictive of the strength and persistence of drug associations was unclear.

Previous work suggests that differences across individuals in phasic dopamine after drug exposure are predictive of drug seeking behavior. For example, mice with higher escalations of cocaine self-administration over a long-access paradigm show a correlated decrease in the phasic dopamine response to drug infusion (Willuhn et al., 2014), and mice who drink more ethanol show less phasic activity in VTA dopamine neurons than do low-drinking mice (Juarez et al., 2017).

Our work suggests that ACh variation may also be important in determining individual differences in drug associations, as we observed that individuals with lower ACh event rates have longer lasting cocaine-context associations. This is consistent with the effects of manipulating ChINs, and suggests that ACh demarcates susceptibility to the influence of context in motivating drug-related behaviors. While further work is needed, this could imply that individuals with ChIN circuitry that is more dysregulated following drug taking may struggle to extinguish drug-context associations, making them vulnerable to future context-induced drug taking. As our experiment used a brief drug conditioning paradigm, a remaining question is whether more chronic drug exposure and drug self-administration paradigms would generate longer-lasting individual differences in ACh signaling.

What might underlie the observed variability in ACh event frequencies across mice? There are multiple mechanisms that have been described by which drug exposure or dopamine signaling can influence ChIN activity. For one, a recent study showed that mice susceptible to cocaine seeking exhibited higher expression of inhibitory D2Rs on ChINs themselves, implicating dopamine action directly on ChINs as a key factor in addiction susceptibility (Lee et al., 2020). Additionally, it has been shown that stimulation of dopamine terminals onto NAc ChINs causes a burst response, and a single dose of amphetamine reduces this burst response by attenuating glutamate co-release (Chuhma et al., 2014), suggesting that excitatory inputs onto ChINs undergo rapid drug modifications that may affect phasic ACh signaling. Recent work has also demonstrated the importance of GABAergic projections from ventral tegmental area onto ventral NAc ChINs in promoting reward reinforcement behavior (Al-Hasani et al., 2021), though whether these projections undergo drug-related plasticity is unknown. Lastly, polysynaptic inhibition can promote local synchronized activity between ChINs, and dopamine can attenuate this inhibition via D2Rs (Dorst et al., 2020). Going forward, *in vivo* examination of the interplay between ACh, dopamine, and other neurotransmitter action with high temporal resolution afforded by new biosensors may clarify the relationships between these signals and reveal modulatory patterns that are predictive of stronger, more persistent addictive behaviors.

The ACh events we detect *in vivo* are likely the product of coordinated spike responses between multiple ChINs. Neighboring ChINs are known to exhibit highly synchronous activity (Ding et al., 2010; Howe et al., 2019), and synchronized burst events are believed to be important for modulating NAc circuit activity and plasticity, causing both local release of dopamine (Cachope et al., 2012; Threlfell et al., 2012) and suppression of MSN firing (English et al., 2011; Witten et al., 2010). The fact that the recorded ACh events likely represent coordinated bursting may explain why we observed decreases in this measure during acute cocaine, while ChIN firing rates are thought to increase during acute cocaine exposure (Berlanga et al., 2003; Lee et al., 2020; Witten et al., 2010).

### Weakening of glutamatergic inputs in NAc across MSN subtypes as a key component of ACh-mediated extinction

Activation of ChINs during a single extinction session decreased cocaine preference and produced similar synaptic plasticity to that observed after multiple extinction sessions, suggesting that ChIN activation serves to accelerate extinction through the same mechanisms that occur endogenously. Specifically, both manipulations produced lower mEPSC frequencies at both D1R and D2R MSNs (**Figures 1 & 3**). Taken together with the relationship between ACh levels in NAc and individual differences in drug extinction (**Figure 2**), these results support the idea that weakening glutamatergic inputs onto both MSN subtypes is a key extinction-related process and that endogenous ACh contributes to this change during repeat extinction.

Our observation of similar plasticity during cocaine extinction on both MSN subtypes is somewhat surprising in light of the distinct and sometimes opposing contributions the two populations have on reward-related behaviors (Bock et al., 2013; Cole et al., 2018; Durieux et al., 2009; Gallo et al., 2018; Hikida et al., 2010; Kravitz et al., 2012; Lobo et al., 2010; O’Neal et al., 2020). What might be the functional significance of this observation of similar plasticity on both MSN subtypes? One possibility relates to the fact that activity in both MSN subtypes are thought to change with cocaine CPP: D1R MSNs show increased activity prior to entry into the cocaine zone (Calipari et al., 2016), and D2R MSNs preferentially fire in the cocaine zone compared to the saline zone (Sjulson et al., 2018). Thus, it may be that extinction requires weakening of inputs at D1R MSNs to decrease likelihood to enter the cocaine zone, and weakening inputs at D2 MSNs to decrease the likelihood that the animal remains in the cocaine zone. Mechanistically, the reduction of presynaptic glutamatergic strength nonspecifically at MSNs may relate to *in vitro* experiments demonstrating that acute muscarinic activation inhibits glutamatergic afferents across MSN subtypes (Ding et al., 2010) (Barral et al., 1999; Malenka and Kocsis, 1988; Pakhotin and Bracci, 2007) via action at the M2- and M3-type muscarinic receptors on these afferents (Hernández-Echeagaray et al., 1998; Higley et al., 2009; Sugita et al., 1991).

In addition to presynaptic weakening across MSN subtypes, we also observed a D2R MSN-specific postsynaptic weakening following ChIN activation that was not evident in the repeat extinction experiments. This difference may be because the optogenetic stimulation generates ACh levels not reached endogenously. It may also be that ChIN-mediated changes in AMPA receptor expression at D2R MSNs during the multiple days of our repeat extinction paradigm are occluded by other factors affecting AMPA receptor expression. For example, it is known that AMPA receptor expression is upregulated by repeat cocaine administration, but that these changes are generally only detectable days after the final cocaine treatment (Kourrich et al., 2007).

## Conclusions

Together, these data point to a model in which phasic ACh promotes cocaine-context extinction by driving a non-specific weakening of glutamatergic inputs across MSN subtypes, with individual differences in the rate of these phasic ACh events controlling the rate of extinction. Thus, individual differences in phasic ACh release may mark a point of susceptibility, as individuals with reduced ACh signaling following drug exposure may be more behaviorally influenced by drug- associated contexts.

## Methods

### Mice

For plasticity experiments, ChAT::IRES-Cre mice (JAX stock 006410: B6;129S6- Chattm2(cre)Lowl/J [RRID: IMSR_JAX:006410]) were maintained on a C57/BL6J background. We used male mice resulting from the cross of ChAT::IRES-Cre mice and Drd1a-tdTomato mice (JAX stock 016204: B6.Cg-Tg(Drd1a-tdTomato)6Calak/J [RRID:IMSR_JAX:016204]). We also used male mice resulting from the cross of ChAT::IRES-Cre mice and Drd2-EGFP mice (MMRC stock To ensure these mice were on a C57/BL6J background, Drd2-EGFP had been backcrossed with C57/BL6J mice for at least four generations.

To measure acetylcholine biosensor fluorescence in NAc, we used ChAT::IRES-Cre mice from the above breedings that either did not express a BAC transgene for fluorophore expression or expressed Drd1a-tdTomato.

Mice were of ages 9-20 weeks. Mice were group-housed with 2–5 mice/cage on a 12-h on, 12-h off light schedule. All behavioral testing was performed during the light off time. All experimental and surgical protocols were approved by Princeton University IACUC to meet guidelines of the NIH guide for the Care and Use of Laboratory Animals.

### Cocaine conditioned place preference

On the first day, each mouse was placed on the left side of the CPP chamber allowed to freely explore the entire apparatus for 15 min (pre-test). Conditioning occurred on days 2 and 3. On each conditioning day, each mouse was confined to one of the side chambers for approximately 18-20 min in the morning and then to the opposite chamber in the afternoon for the same period of time. Before placement into a given chamber, intraperitoneal injections of cocaine (15 mg/kg) before placement in one chamber or intraperitoneal injections of an equal volume of saline (∼0.1 mL) before being placed in the other chamber. On day 4, extinction testing began. On this day, mice were allowed to again freely explore the entire apparatus for 15 min.

For the repeat extinction experiment (**Figure 1**), one group of mice underwent an extinction test each day from days 4-7. Immediately following the final extinction test, mice were killed for *ex vivo* electrophysiology. Another group of mice underwent conditioning, but never returned to the behavior room after their final conditioning session. These mice were also killed on day 7 for *ex vivo* electrophysiology at approximately the same time of day as mice who underwent repeat extinction testing (1200 hours).

For the acetylcholine sensor experiment (**Figures 2-3**), before each session, mice were connected to the photometry patch cable and placed in their home cage for 10 mins. After the session, mice were returned to their home cage for 5 mins. Mice remained plugged into the patch cable during injections for conditioning sessions.

For the optogenetic experiment (**Figure 4**), on the day 1 Baseline session, mice were connected to patch cables that were not emitting light. On the day 4 extinction test, mice were again connected to the patch cables. Experimental mice received optogenetic stimulation during the 15 min extinction test. All mice were killed immediately afterward for *ex vivo* electrophysiology. See **Optogenetic stimulation of cholinergic interneurons** for additional details.

### Recordings of acetylcholine biosensor fluorescence

Mice were injected unilaterally or bilaterally with 600 nL of AAV 2/9 with hSyn-GACh3.0-WPRE-hGHpA (Jing et al., 2020) (PNI Viral Core Facility, injected titer of 6.5*10^13^ genome copies/mL) in the ventral-medial NAc in two sites (M-L, 0.65 mm; A-P, 1.43 mm; D-V, 4.75 mm) and (M-L, 0.65 mm; A-P, 1.43 mm; D-V, 4.55 mm). Optical fibers (400 μm core diameter, 0.48 NA) were implanted unilaterally or bilaterally at a 10 degree angle to target the injection region in the NAc (non-rotated coordinates: M-L, 0.685 mm; A-P, 1.43 mm; D-V, 4.55 mm; rotated coordinates at 10 degrees: M-L, 1.46 mm; A-P, 1.43 mm; D-V, 4.36 mm).

To record neural activity with GACh3.0 expressed non-specifically in NAc tissue, mice were connected to a fiber photometry setup (Cai et al., 2020; Parker et al., 2016). In all cases, recordings were unilateral from the same site in a given mouse; in bilaterally surgerized mice, the recording site was selected based on the apparent quality of the fluorescence signal at least one day before behavior testing. Light from the excitation laser (488 nm; Micron Technology) was filtered (FL488, Thorlabs) then passed through a dichroic mirror (MD498, Thorlabs) and traveled through a patch cable (Mono Fiber-optic Patchcord, 400 µm core, 0.48 NA, Doric Lenses) coupled via ceramic split sleeve (2.5 mm diameter, Precision Fiber Products) to the optic fiber implanted in the mouse brain. Laser light delivery was controlled by a lock-in amplifier (Ametek, 7265 Dual Phase DSP Lock-in Amplifier), which delivered light at 210.999 Hz, and the light intensity at the tip of the patch cable was approximately 20 µW. Fluorescent emission from GACh3.0 was then passed through the same patch cable, filtered (MF525-39, Thorlabs), and passed through the same dichroic mirror into a photodetector (Model 2151, New Focus), and the signal was acquired using the same-lock-in amplifier, with a time constant of 20 ms. AC gain on the lock-in amplifier was set to 0 dB. Signal was digitized at 100 Hz on a data acquisition board (USB-201, Measurement Computing) and stored.

dF/F was calculated by the following formula:

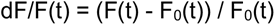

Where F0(t) is an estimate of the baseline fluorescence at time t. To isolate transient events from slow fluctuations in the signal, F0(t) calculated as the fifth percentile based on the preceding 5 s of the recording. Z-score of dF/F signal was calculated across experiment days for each mouse; the mean and standard deviation used for Z-score were the mean and standard deviation of the concatenated dF/F signal across all behavioral sessions from all 7 experiment days.

For event detection from the DF/F, for each mouse, we determined a threshold amplitude that local maxima in the z-scored trace must surpass. To calculate this threshold, we used a heuristic on the data after omitting large peaks. To omit large peaks, we first calculated an initial threshold which was set to 2 * the mean absolute deviation (MAD) above the median of the z-scored trace. We used the MATLAB function “findpeaks” to find peaks with amplitudes that exceeded this initial threshold, and omit indices +/- half of the width at half max. The threshold for peak detection was set to 2*MAD above the median of the trace with large peaks excluded. Peak detection was then performed on the original z-scored traces using the MATLAB function “findpeaks” with the minimum peak height set to the threshold, and a minimum inter-peak-interval of 100 ms, meaning that if there were multiple events within 100 ms, only the 1st was recorded.

### Optogenetic stimulation of cholinergic interneurons

During surgery, AAV5-EFIa-DIO-ChR2-eYFP (Princeton Viral Core, injected 1 uL per hemisphere at titer of 1.2 x 10^14^) was infused bilaterally in ventral medial NAc (M-L, +/- 0.65 mm; A-P, 1.43 mm; D-V, 4.75 mm). Optic fibers (300 mm core diameter, 0.37 NA) delivering 10 mW of 447 nm laser light (measured at the end of the patch cable) were implanted bilaterally at a 10 degree angle to target the ventral medial NAc (non-rotated coordinates: M-L, +/- 0.77 mm; A-P, 1.43 mm; D-V, -4.45 mm; rotated coordinates at 10 degrees: M-L, +/- 1.55 mm; A-P, 1.43 mm; D-V, -4.25 mm). Animals were anesthetized for implant surgeries with isoflurane (3–4% induction and 1–2% maintenance). Behavioral testing occurred 3-4 weeks after surgery to allow for animal recovery and viral expression. Mice were plugged into the patch cables during the baseline and the extinction test sessions. In experimental animals, during extinction testing, light was administered throughout the test in a burst pattern with 5 ms wide pulses at 15 Hz for 2s interleaved with 2s light off periods. Control animals received the same surgery and were plugged into the same patch cable but no light was administered during extinction testing.

### *Ex vivo* electrophysiology

Mice were anesthetized with an i.p. injection of Euthasol (0.06ml/30g). Mice were decapitated and the brain was extracted. After extraction, the brain was immersed in ice-cold NMDG ACSF (92 mM NMDG, 2.5 mM KCl, 1.25 mM NaH2PO4, 30 mM NaHCO3, 20 mM HEPES, 25 mM glucose, 2 mM thiourea, 5 mM Na-ascorbate, 3 mM Na-pyruvate, 0.5 mM CaCl2·4H2O, 10 mM MgSO4·7H2O, and 12 mM N-Acetyl-L-cysteine; pH adjusted to 7.3-7.4) for 2 minutes. Afterwards coronal slices (300 µm) were sectioned using a vibratome (VT1200s, Leica, Germany) and then incubated in NMDG ACSF at 34 C for approximately 14 minutes. Slices were then transferred into a holding solution of HEPES ACSF (92 mM NaCl, 2.5 mM KCl, 1.25 mM NaH2PO4, 30 mM NaHCO3, 20 mM HEPES, 25 mM glucose, 2 mM thiourea, 5 mM Na-ascorbate, 3 mM Na pyruvate, 2 mM CaCl2·4H2O, 2 mM MgSO4·7H2O and 12 mM N-Acetyl-l-cysteine, bubbled at room temperature with 95% O2/ 5% CO2) for at least 45 mins until recordings were performed.

Whole-cell recordings were performed using a Multiclamp 700B (Molecular Devices, Sunnyvale, CA) using pipettes with a resistance of 4-7 MOhm filled with an internal solution containing 100 mM cesium gluconate, 0.6 mM EGTA, 10 mM HEPES, 5 mM NaCl, 20 mM TEA, 4 mM Mg- ATP, 0.3 mM Na-GTP and 3 mM QX 314 with the pH adjusted to 7.2 with CsOH and the osmolarity adjusted to around 289 mmol kg−1 with sucrose. During recordings, slices were perfused with a recording ACSF solution (120 mM NaCl, 3.5 mM KCl, 1.25 mM NaH2PO4, 26 mM NaHCO3, 1.3 mM MgCl2, 2 mM CaCl2 and 11 mM D-(+)-glucose, and was continuously bubbled with 95% O2/5% CO2). Infrared differential interference contrast–enhanced visual guidance was used to select neurons that were 3–4 cell layers below the surface of the slices. MSNs were identified by the presence of either eGFP or td-Tomato of BAC transgenic mice using a fluorescence microscope (Scientifica SliceScope Pro 1000; LED: SPECTRA X light engine (Lumencor)). The recording solution was delivered to slices via superfusion driven by peristaltic pump (flow rate of 4-5 ml/min) and was held at room temperature. The neurons were held at −70 mV (voltage clamp), and the pipette series resistance was monitored throughout the experiments by hyperpolarizing steps of -10 mV with each sweep. If the series resistance changed by >20 % during the recording, the data were discarded. Whole-cell currents were filtered at 1 kHz and digitized and stored at 10 KHz (Clampex 10; MDS Analytical Technologies). All experiments were completed within 4 hours after slices were made to maximize cell viability and consistency.

Miniature EPSCs (mEPSCs) were recorded in the presence of TTX (1 μM), d-AP5 (50 μM) and picrotoxin (100 μM) in the recording ACSF solution. mEPSC data was analyzed with Stimfit (Guzman et al., 2014) using detection threshold of >7 pA and rise time <3 ms, and the results were visually verified. For each cell, a stretch of 300 mEPSCs were analyzed, with data collection beginning 10-15 minutes after patching onto each cell.

For paired pulse ratio (PPR) and AMPA/NMDA ratio measurements, picrotoxin (100 μM) was added to the recording ACSF solution. A bipolar stimulating electrode was placed at the medial-ventral edge of the NAc. For PPR, every 30 s, a paired-pulse stimulation was delivered with a 50 ms inter-stimulus interval (ISI). Stimulation duration was 0.1 ms and current was approximately 0.02-0.08 mA. Evoked EPSCs were recorded from MSNs at a holding potential of -70 mV. 10 repetitions of the stimulation protocol were recorded per cell after stable evoked EPSCs were achieved. PPR was calculated as the ratio between the peak amplitudes of the second and first EPSC. To determine AMPA receptor and NMDA receptor currents, evoked EPSCs were recorded from MSNs at a holding potential of +40 mV. After approximately 10 stimulations, d-AP5 (50 μM) was bath applied to block NMDA receptor currents and isolate AMPA receptor currents. At least 10 evoked EPSCs were recorded following bath application of dAP-5. The NMDA receptor current was calculated as the difference of the mean currents in the absence and presence of d-AP5. In example traces, stimulation artefacts have been removed.

For temporal summation and intrinsic excitability experiments, pipettes were filled with a potassium-based internal solution containing 120 mM potassium gluconate, 0.2 mM EGTA, 10 mM HEPES, 5 mM NaCl, 1 mM MgCl2, 2 mM Mg-ATP and 0.3 mM NA-GTP, with the pH adjusted to 7.2 with KOH.

For the temporal summation experiment, the internal solution also contained 50 μM Alexa Fluor 594 hydrazide for visualization of dendrites. EPSP-like depolarizations mediated by activation of AMPA receptors were evoked using short iontophoretic pulses (10 pulses at 20 Hz with duration 3 ms at 200 nA) of sodium glutamate (150 mM) (Müller and Remy, 2013; Shen et al., 2007) from a puff pipette (pipette resistance 30 MOhm) to a region of visibly identified dendrite approximately 50-100 µm from the soma. Sodium glutamate solution also contained 50 μM Alexa Fluor 594 hydrazide for visualization of puff pipette tip and confirmation of release. Experiments were performed with TTX (1 μM), d-AP5 (50 μM) and picrotoxin (100 μM) in the recording ACSF solution.

For intrinsic excitability experiments, current-clamp recordings were performed in which the resting membrane potential was normalized to -80 mV. In some cases, the resting membrane potential was brought to -80 mV by small current injections of <50 pA. If the compensating current was >50 pA, the recording of the neuron was terminated (Mu et al., 2010). To measure evoked spiking, a current step protocol (-200 to +350 pA; 50 pA increment; 15 s interpulse interval) was run for 5 runs (Mu et al., 2010).

### *Ex vivo* confirmation of ACh sensor response selectivity

To confirm that tissue expressing GACh3.0 was responsive to acetylcholine but not cocaine, mice were injected bilaterally with 600 nL of AAV 2/9 with hSyn-GACh3.0-WPRE-hGHpA (PNI Viral Core Facility, injected titer of 6.5*10^13^ genome copies/mL) in the ventral-medial NAc in two sites (M-L, 0.65 mm; A-P, 1.43 mm; D-V, 4.75 mm) and (M-L, 0.65 mm; A-P, 1.43 mm; D-V, 4.55 mm). 3 weeks after surgery, we performed *ex vivo* slice imaging. Brain slices were collected using the methods and solutions described in ***Ex vivo* electrophysiology**. During recordings, slices were perfused with the recording ACSF solution (see ***Ex vivo* electrophysiology**). Fluorescence was imaged using a CMOS camera (ORCA-Flash 2.8, Hamamatsu) at 33.333 Hz (30 ms exposure windows) using a GFP filter cube set (excited ET470/40x, dichroic T495LP, emitter ET525/50m) (Fleming et al., 2021). To test for fluorescence responses to acetylcholine or cocaine, a glass pipette filled with recording ACSF containing a given drug (100 µM acetylcholine, 10 µM cocaine, or 100 µM cocaine) was placed above a segment of tissue. Slight positive pressure (approximately 80 kPa) was briefly applied (100 ms), and time-locked fluorescence responses were recorded. Recordings were performed on the same segment of tissue, but for each drug the pipette was replaced and repositioned over the tissue.

### Immunohistochemistry

Mice were first anesthetized and then transcardially perfused with cold 4% paraformaldehyde (PFA) in PBS (pH 7.4). Brains were fixed in 4% PFA. 50 μm coronal slices were sectioned on a vibratome, and then were subsequently stored in cryoprotectant at 4 °C. For immunohistochemistry, individual sections were washed in PBS and then incubated for 30 min in 0.3% Triton-X and 3% normal donkey serum (NDS).

For ChAT labeling, ChAT primary antibody (1:200; Millipore, product# AB144P [RRID:AB_2079751]) incubations were performed overnight at 4 °C in 3% NDS/PBS. Sections were washed and left to incubate in secondary antibodies conjugated to AlexaFluor586 for 3 hrs at room temperature (1:1000; Life Technologies, Product# A11057 [RRID:AB_10564097]).

For enhancement of GFP in GACh3.0 tissue labeling, GFP primary antibody (1:500; Millipore, product # NB600-308 [RRID:AB_10003058]) incubations were performed overnight at 4 °C in 3% NDS/PBS. Sections were washed and left to incubate in secondary antibodies conjugated to AlexaFluor586 for 3 hrs at room temperature (1:1000; Life Technologies, Product# A11057 [RRID:AB_10564097]).

Following a 20 min incubation with DAPI (1:50,000) sections were washed and mounted on microscope slides with Fluorogold (Southern Biotechnology, product # 0100-01).

### In situ RNA hybridization

For the in situ RNA hybridization experiments, mice were anesthetized with 0.1 mL Euthasol (i.p. injection) and transcardially perfused with 4% PFA in PBS. Brains were dissected out and post-fixed in 4% PFA overnight. Brains were then put through a sucrose gradient: 10% sucrose in PBS solution for 6-8 hours, 20% sucrose in PBS solution overnight, and then 30% sucrose in PBS solution overnight. 18 µm thick coronal sections containing the nucleus accumbens were cut on a cryostat. In situ hybridization was performed on mounted sections using the RNAscope Multiplex Fluorescent Assay (Advanced Cell Diagnostics, Inc., No. 323110). Custom probes were used for Mm-Drd1a-C1 (406491) and Mm-Drd2-C2 (406501-C2, 1:50 dilution in C1 probe solution), and Mm-tdTomato-C3 (317041-C3, 1:50 dilution in C1 probe solution for drd1a- tdTomato mice only). The following fluorophores (Perkin Elmer, NEL760001KT) were used to report RNA detection in each channel: fluorescein (C2 in drd1a-tdTomato slices), cy3 (C3 in drd1a-tdTomato slices; C1 in drd2-eGFP slices), and cy5 (C1 in drd1a-tdTomato slices; C2 in drd2-eGFP slices). Fluorophores were reconstituted in 60 mL DMSO and diluted in TSA buffer provided in the RNAscope kit at a concentration of 1:1200. Following the in situ hybridization protocol, a GFP antibody stain was used to enhance visualization of drd2-eGFP expression. The primary antibody was a mouse monoclonal anti-GFP (1:1000 dilution, Life Technologies, No. G10362) and the secondary was a donkey anti-rabbit coupled to Alexa 488 (1:1000 dilution, Jackson ImmunoResearch, No. 711-545-152). Cellular resolution images of mounted coronal sections were acquired using a Zeiss confocal microscope (LSM 700; four visible solid-state lasers: UV 405; argon 458/488; HeNe 555/568; far-red 639) (Oberkochen, Germany). The first example image in **Figure 2a** was taken at 63x, 0.7x zoom with oil immersion, and the second image was taken at 63x, 2.0x zoom with oil immersion.

### Statistical analyses

Behavioral preference for repeat extinction CPP experiments with only one experimental group undergoing repeat preference (**Figure 1b, Supplementary figure 5a**) was assessed using the “aov” function in R to perform a one-way repeated measures ANOVA with session as fixed effect and mouse as random effect. Post-hoc tests comparing Baseline to Test 1 preference were performed using the “pairwise.t.test” function in R to perform a pairwise t-test between preference during the two sessions with Holm’s adjustment for multiple comparisons.

To compare changes in ACh event frequency in **Figures 2f-g**, we used pairwise t-tests, using the “ttest” function in MATLAB, of mean event frequency between Baseline and Test 1 for the 10 min period before behavior, the 15 min behavioral period, and the 5 min period after behavior. To compare event frequency based on chamber side, pairwise t-tests were performed using the mean frequencies for each mouse on the saline vs cocaine zone. To measure the relationship between event frequency on Test 1 and preference on Tests 1-4 (**Figure 2j, Supplementary figure 5c-f**), we calculated the correlation coefficient and corresponding p values between Test 1 frequency and test preferences using the “corrcoef” function in MATLAB. Additionally, for **Figure 2k**, we measured the relationship between Test 1 frequency groups and preference on Tests 1-4 with a repeated measures ANOVA using the “aov” function in R, where preference was predicted using median split frequency group, session (Tests 1-4), and their interaction as fixed effects, and mouse as a random effect.

To measure how speed was affected by ACh events (**Figure 3b-d**), we calculated the baseline-corrected change in speed around the time of each event. Baseline speed was calculated as mean speed in a 1 s window starting 2 s before the ACh event peak time. The speed around each event was calculated as the mean speed in a 1.5 s window beginning 0.5 s before the time event peak time and ending 1 s after the event peak time. The change in speed was calculated as the difference between the baseline speed and the speed around the time of the event (**Figure 3d**).

To measure the relationship between speed and event frequency (**Figure 3e**), 10 speed deciles were first calculated for each mouse using combined speed data from both the Baseline and Test 1 sessions. For each session, the number of events occurring while the animal was moving at a speed within a given decile range was counted and divided by the time the animal spent in that speed range (1/10 of total time between the two behaviors) to calculate event frequencies. Mean event frequencies were calculated for each speed decile for Baseline and Test 1. To quantify the relationship between speed and event frequency, we performed a linear, mixed-effects regression using the “lmer” function in R, with frequency predicted using speed decile, behavioral session, and their interaction as fixed effects, and mouse as a random effect. Effect size was calculated by inputting the resulting model into the “lme.dscore” function in the “EMAtools” package in R. To compare how the slope of the speed decile vs. event frequency plots corresponded to preference on Tests 1-4 (**Figure 3f**), slopes for each mouse were first calculated by fitting the mean event frequency across speed deciles by fitting the data to a linear function using the “polyfit” function in MATLAB. Slopes were median split, and we performed a repeated measures ANOVA using the “aov” function in R, where preference was predicted using the median split group, session (Tests 1-4), and their interaction as fixed effects, and mouse as a random effect.

To compare differences in preference between experimental groups in the optogenetic experiment (**Figure 4e; Supplementary figure 12a-c**), we performed a multi-factor, repeated measures ANOVA using the “aov” function in R, where preference was predicted using mouse transgenic line (drd1a-tdTomato or drd2-GFP), group (Control vs ChIN activation), and session (Baseline vs Test), and their interactions as fixed effects, with mouse as a random effect. Two post-hoc t-tests were performed between groups (Control vs ChIN activation) for both Baseline and Test days using the “pairwise.t.test” function in R. To compare differences between groups (Control vs ChIN activation) as a function of time during the Test session (**Figure 4f**), we first calculated the fraction of time each animal spent in the cocaine zone during the Test, divided into unsmoothed, non-overlapping 1 minute bins. With this data, we used the “lmer” function in R to perform a linear, mixed-effects regression on preference for the cocaine zone using group (Control vs ChIN activation), time (minute bins 1-15), and their interaction as fixed effects, with mouse as a random effect. We performed a series of post-hoc t-tests using the “pairwise.t.test” function, with Holm’s correction, to determine at which time points were the Control and ChIN activation groups different.

To compare the mEPSC frequency and amplitudes across conditions, linear, mixed-effects regressions were performed using the “lmer” function in R. Interevent interval (IEI) data was log-transformed while amplitude data was inverse-transformed to best fit a normal distribution. For statistical analysis of both IEI data and amplitude data for **Figures 1e-h**, the linear, mixed effects regression model included group (No extinction vs. Repeat extinction) as a fixed effect and cell as a random effect. For statistical analysis of both IEI data and amplitude data for **Figures 4h-k**, the linear, mixed effects regression model included group (Control vs. ChIN activation) as a fixed effect and cell as a random effect. Effect sizes were calculated as Cohen’s d using the “lme.dscore” function in the “EMATools” package in R. Separate regressions were calculated for data from D1R and D2R MSNs.

To compare differences in paired pulse ratio and AMPA/NMDA ratio between groups (Control vs ChIN activation; **Figure 4l-q**), we performed two-tailed t-tests using the “ttest2” function in MATLAB. Effect sizes were calculated as Cohen’s d using custom written software in MATLAB, where the difference in means between the two data sets is divided by the pooled standard deviations of the two datasets.

To compare the difference in temporal summation of excitatory EPSPs in the glutamate iontophoresis experiment (**Figure 4r-s**), for each recording all EPSPs were first normalized to the amplitude of the first EPSP in the respective trace. We compared the amplitudes of these normalized EPSPs between groups (Control vs ChIN activation) by using the “lmer” function in R to perform a linear, mixed effects regression on EPSP amplitude using group, EPSP number, and their interaction as another fixed effect, with cell as a random effect.

To compare the differences in evoked spikes as a function of injected current between groups (**Figure 4u**), we first calculated the mean number of spikes for each current step across all runs for a given cell. If the mean number of spikes decreased with increasing current injection (because of depolarization block), data from that current step and above was removed. Separately for D1R and D2R MSN data, we used the “lmer” function in R to perform a linear, mixed-effects regression on the number of evoked spikes, using group (Control vs ChIN activation), injected current value, and their interaction as fixed effects, with cell as a random effect. We also inputted the resulting model into R function “testInteractions” to determine whether the slopes of the current vs. evoked spikes lines were significantly different between groups (Control vs ChIN activation). Effect sizes were calculated using the “lme.dscore” function in the “EMAtools” package in R.

For action potential statistics in the evoked spiking experiment (**Figure 4t-u; Supplementary figure 14**), only the first spike in each run was used. Action potential threshold was calculated as the first point of positive acceleration of voltage ((ΔV/Δt) / Δt) that exceeded 3*SD of membrane noise in the period prior to current injection (Baufreton et al., 2005; Willett et al., 2019). The fast afterhyperpolarization potential was calculated as the voltage value 8 ms following onset of the first actional potential in a run; the medium afterhyperpolarization potential was calculated as the voltage 16 ms following the onset of the first action potential.

## Acknowledgements

We thank V. Corbit and A. Pan-Vazquez for comments on the manuscript, and members of the Witten laboratory for their support. We thank B. Juarez for discussion on individual differences in neuromodulation. We thank E. Engel for reagents. Funding was from NYSCF (IBW); NSF GRFP (WF); ARO W911NF1710554 (IW); U19 NS104648-01 (IBW); and a Simons Investigator Award in Mathematical Modeling of Living Systems (DMW). IBW is a New York Stem Cell Foundation— Robertson Investigator.

## Author contributions

Conceptualization, W.F., J.L., I.B.W.; Methodology, W.F., J.L., B.B, S.B., I.B.W.; Validation, W.F., J.L., B.B., S.B.; Investigation, W.F.; Formal Analysis, W.F., J.L.; Writing - Original Draft, W.F., I.B.W.; Writing - Review & Editing, W.F., J.L., B.B., S.B., I.B.W.; Visualization, W.F.; Resources, I.B.W.; Supervision, J.L., I.B.W. Funding Acquisition, W.F., I.B.W.

## Declaration of interests

The authors declare no competing interests.

## Supplementary figures

**Supplementary figure 1.**
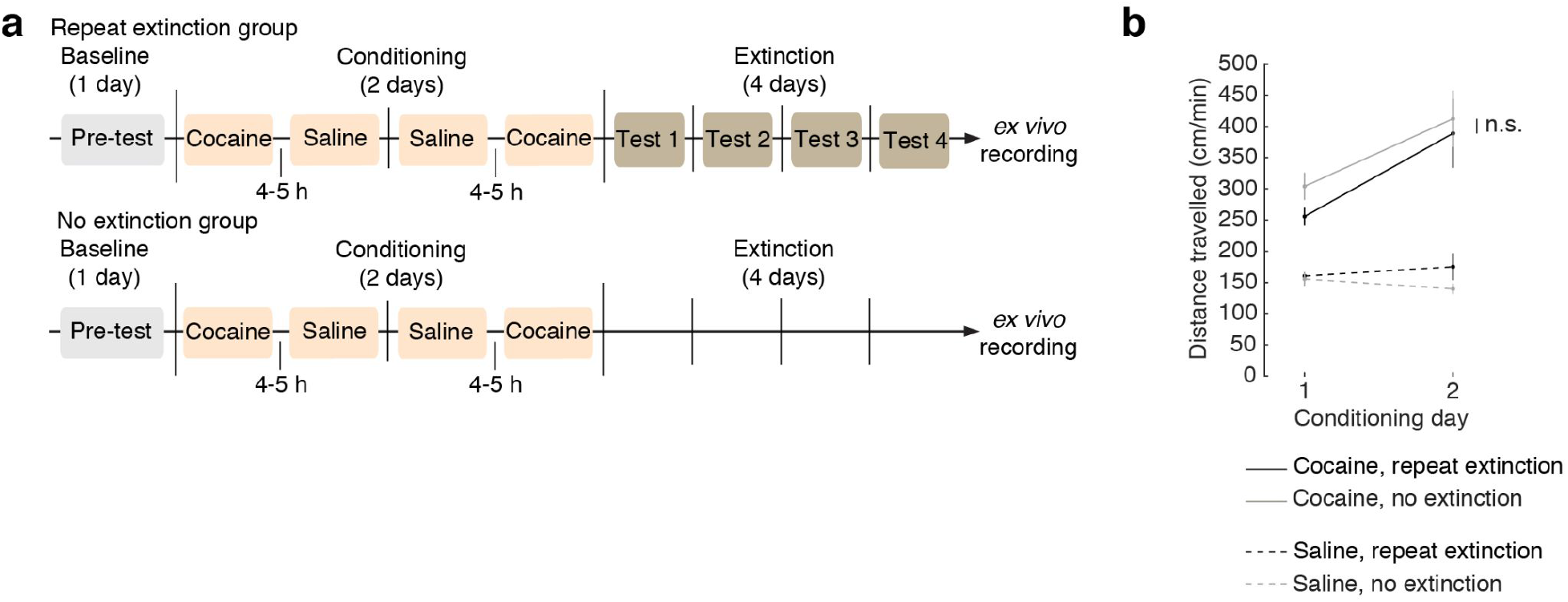
Locomotion during acute cocaine was not different between repeat extinction and no extinction groups, suggesting acute cocaine effects were similar. **a.** Timeline of the CPP experiment from Figure 5, where mice are conditioned with saline and cocaine, followed by either repeat extinction testing or no extinction. **b.** Mean total distance travelled during conditioning sessions with cocaine (solid lines) and saline (dashed lines) for both repeat extinction (black lines) and no extinction (light grey lines) groups. There was no significant difference in total distance travelled during either cocaine session between repeat extinction and no extinction groups, indicating that cocaine treatment had similar acute effects between the two groups (p = 0.125 for cocaine on day 1; p = 0.747 for cocaine on day 2; two-sample t-tests between repeat extinction and no extinction groups).

**Supplementary figure 2.**
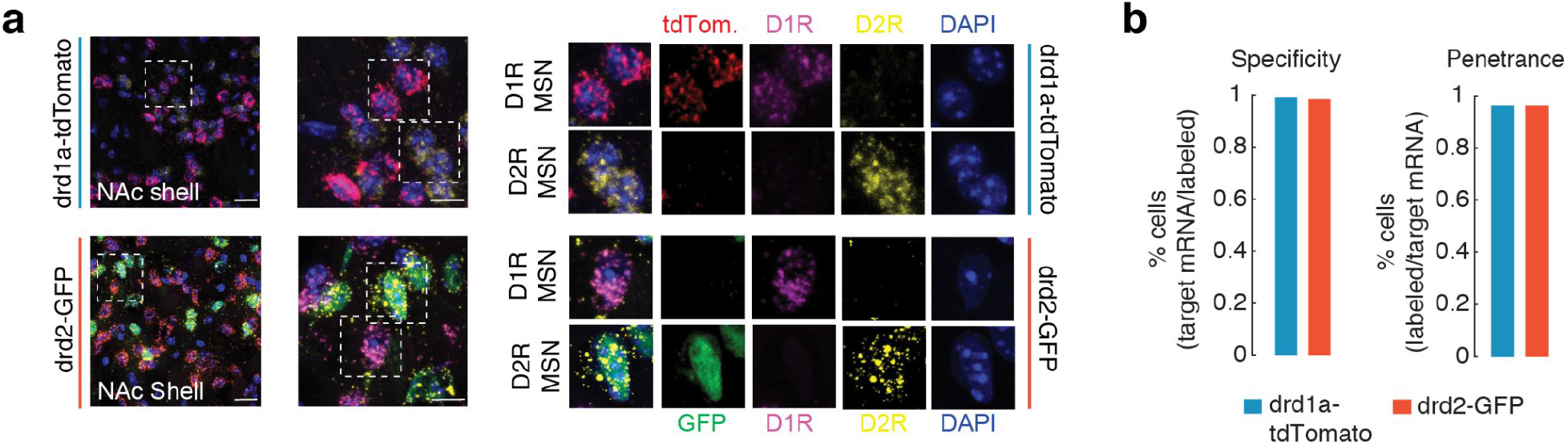
In situ hybridization confirms reliable labeling of MSN subtypes in drd1a- tdTomato and drd2-GFP mice crossed with ChAT-Cre mice. **a.** Example images from neurons in NAc shell showing co-expression of genetically encoded fluorescent protein and appropriate dopamine receptor mRNA. Rows contain representative images from either ChAT-cre::drd1a-tdTomato (top) or ChAT-Cre::drd2-GFP (bottom) tissue. In each row, the scale bar in the first image is 20 µm. The second image is the dotted excerpt from the first image, where the scale bar is 10 µm. Dotted excerpts in the second image highlight either D1R or D2R MSNs. Successive images show fluorescent labeling (tdTomato or GFP) and labeled RNA (D1R and D2R) for a given MSN. **b.** Specificity and penetrance of fluorescent labeling of MSNs in ChAT::Cre/drd1a-tdTomato and ChAT::Cre/drd2-GFP mice. In ChAT::Cre / drd1a-tdTomato mice, tdTomato labeling of D1R was 99.1% specific (of 981 td-Tomato labeled cells, 972 were D1R+), and td-Tomato labeling had 96.6% penetrance (of 1076 D1R+ cells, 1039 were td- Tomato labeled). In ChAT::Cre/drd2-GFP mice, GFP labeling of D2R was 98.7% specific (of 1436 GFP labeled cells, 1417 were D2R+), and GFP labeling had 96.3% penetrance (of 1443 D2R+ cells, 1389 were GFP+).

**Supplementary figure 3.**
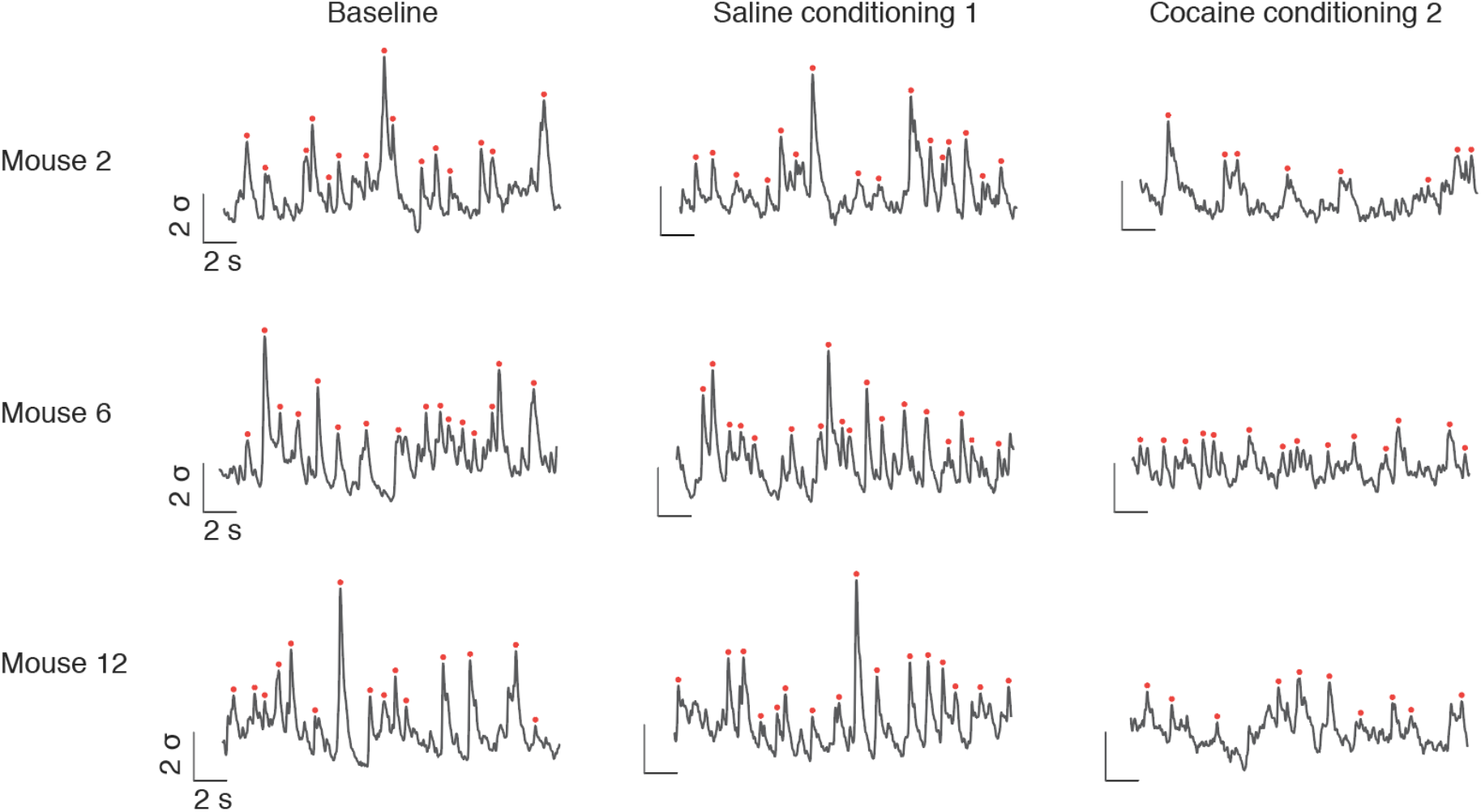
Example event detection on fiber photometry GACh3.0 fluorescence. **a.** Examples of z-scored dF/F of GACh3.0 photometry signal from three mice (rows) across three behavioral sessions (columns; Baseline, Saline conditioning on day 1 of conditioning, Cocaine conditioning on day 2 of conditioning). Signals are z-scores, and significant events are denoted by red circles.

**Supplementary figure 4.**
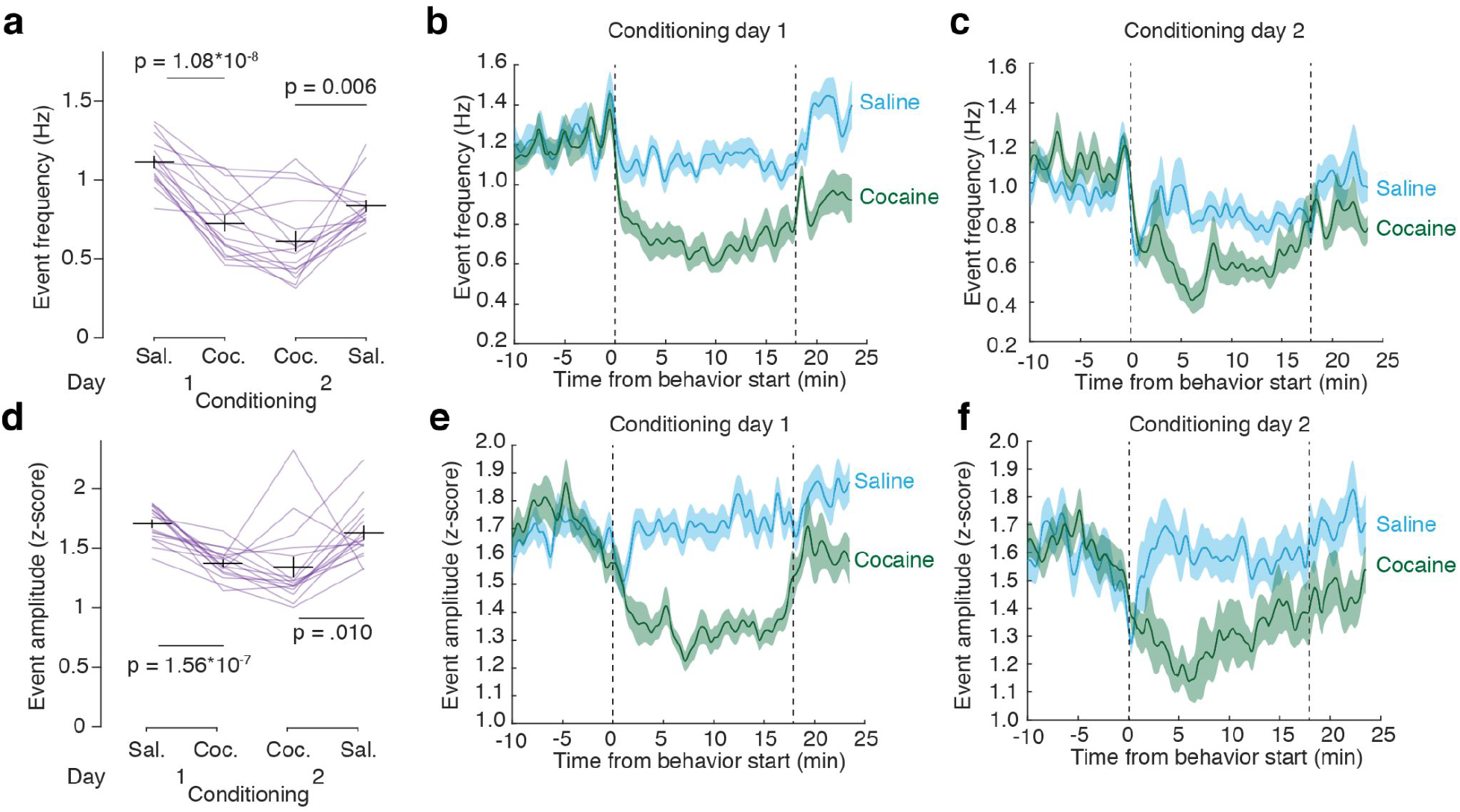
Cocaine conditioning reduces ACh event frequency and amplitude compared to saline conditioning. **a.** ACh event frequency was reduced during cocaine (Coc.) conditioning compared to saline (Sal.) conditioning on days 1 and 2 (p = 1.08*10^-8^ for pairwise t-test between Saline and Cocaine on day 1; p = 0.006 for pairwise t-test between Saline and Cocaine on day 2). **b.** Mean ACh event frequency on the first conditioning day during saline (blue) and cocaine (green) sessions. **c.** Mean ACh event frequency on the second conditioning day during saline (blue) and cocaine (green) sessions. **d.** ACh event amplitude was reduced during cocaine (Coc.) conditioning compared to saline (Sal.) conditioning on days 1 and 2 (p = 1.56*10^-7^ for pairwise t-test between Saline and Cocaine on day 1; p = 0.010 for pairwise t-test between Saline and Cocaine on day 2). **e.** Mean ACh event amplitude on the first conditioning day during saline (blue) and cocaine (green) sessions. **f.** Mean ACh event frequency on the second conditioning day during saline (blue) and cocaine (green) sessions. All shaded regions and vertical bars are SEM.

**Supplementary figure 5.**
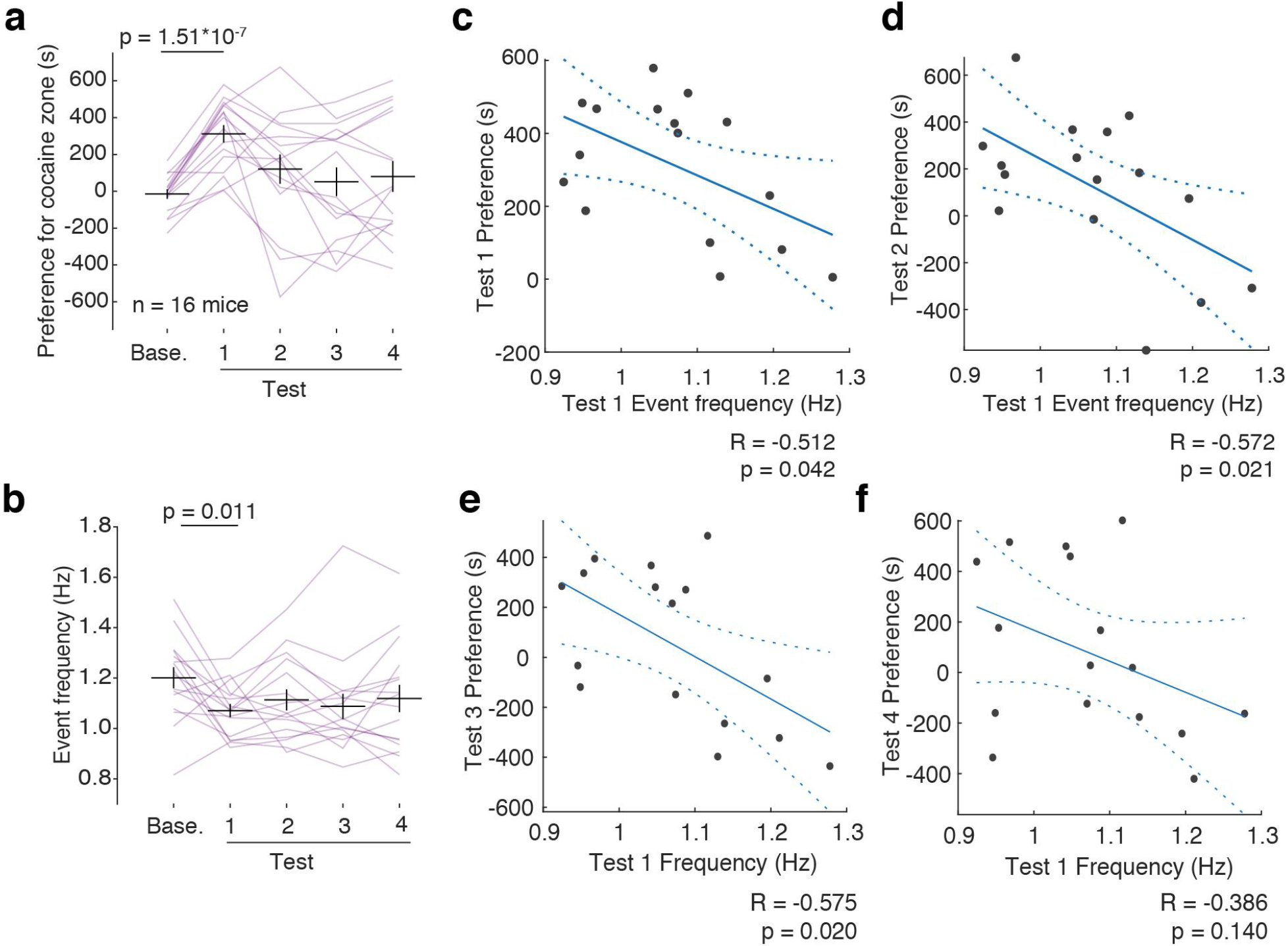
Relationships between mean preference for the cocaine zone and mean event frequency. **a.** Preference for the cocaine zone, measured as the difference in time spent on the cocaine zone minus time spent on the saline zone. Mice from the repeat extinction group preferred the cocaine-paired chamber more on Test 1 than on Baseline, but not on subsequent tests (F(4,60) = 6.801, p = 1.39*10^-4^ for session in one-way repeated measures ANOVA with session as fixed effect and mouse as random effect. Effect size d = 2.297, p = 1.51*10^-7^ for pairwise t-test of Baseline and Test 1, Šidák correction for multiple comparisons). **b.** Mean event frequency during each behavior test session. Mice showed decreased frequency on Test 1 compared to baseline, but not on subsequent tests (F(4,60) = 2.583, p = 0.046 for session in one-way repeated measures ANOVA with session as fixed effect and mouse as random effect. Effect size d = 0.725, p = 0.011 for pairwise t-test of Baseline and Test 1, Šidák correction for multiple comparisons). **c.** Correlation between event frequency on Test 1 and preference for the cocaine zone on Test 1 (correlation coefficient R = -0.512; p = 0.042). **d.** Correlation between event frequency on Test 1 and preference for the cocaine zone on Test 2 (correlation coefficient R = -0.572; p = 0.021). **e.** Correlation between event frequency on Test 1 and preference for the cocaine zone on Test 3 (correlation coefficient R = -0.575; p = 0.020). **f.** Correlation between event frequency on Test 1 and preference for the cocaine zone on Test 4 (correlation coefficient R = -0.386; p = 0.140).

**Supplementary figure 6.**
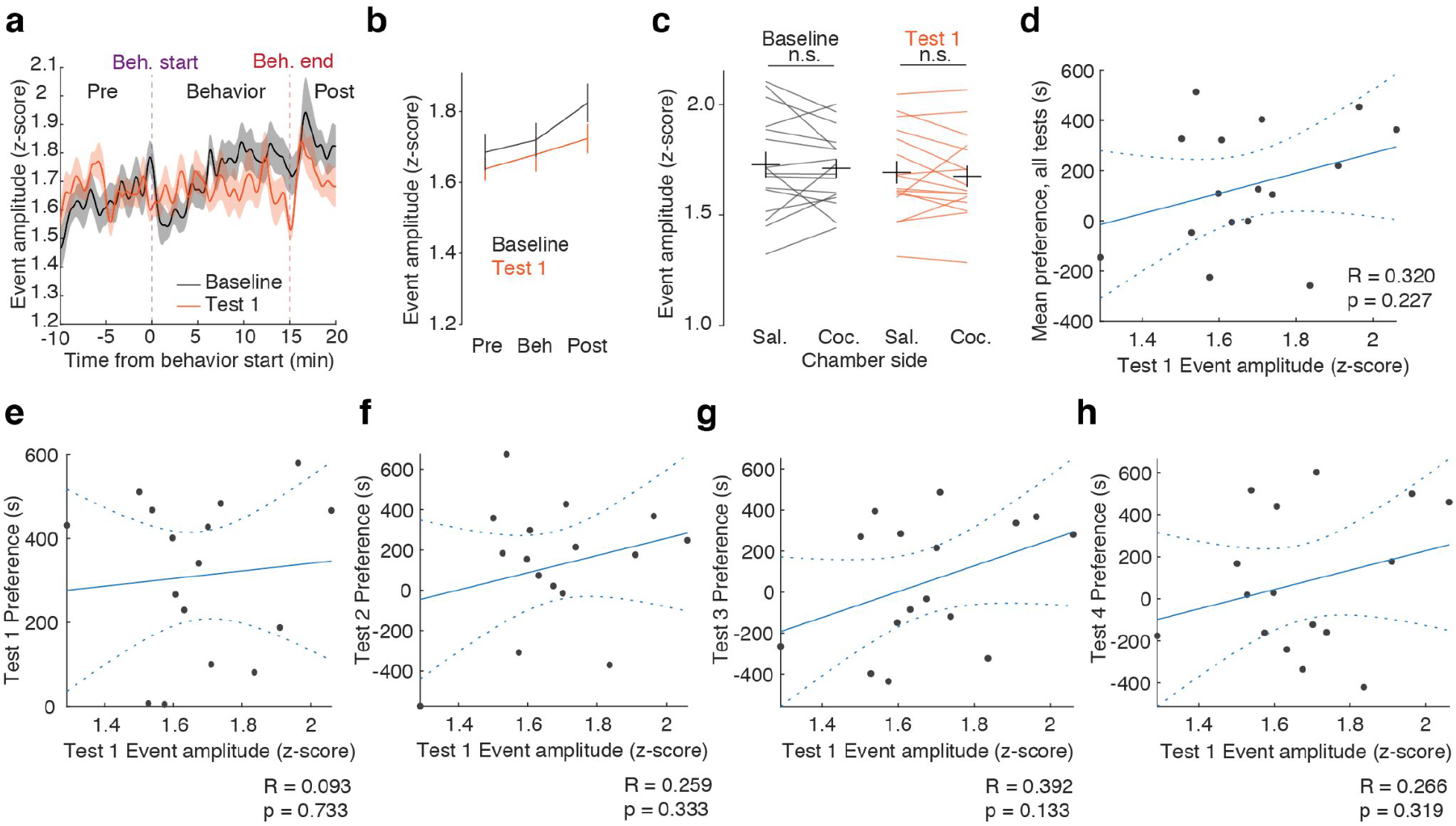
ACh event amplitude is not different from Baseline to Test 1, does not vary by chamber side, and is not related to preference on extinction Tests 1-4. **a.** Mean event amplitude calculated from events within a 30 s moving window before, during, and after Baseline (grey) and Test (1) sessions. **b.** Mean event amplitude is not different between Baseline and Test 1 in pre-behavior (Pre), behavior (Beh), or post-behavior (Post) periods (p = 0.448 for pre-behavior period pairwise t-test of Baseline and Test 1; p = 0.550 for behavior period pairwise t-test of Baseline and Test 1; p = 0.152 for post-behavior period pairwise t-test of Baseline and Test 1). **c.** Mean event amplitude is not different between cocaine- and saline-conditioned zones on Baseline or Test 1 (p = 0.646 for pairwise t-test of Saline and Cocaine zones on Baseline; p = 0.482 for pairwise t-test of Saline and Cocaine zones on Test 1). **d.** Mean event amplitude on Test 1 is not correlated with mean preference for the cocaine zone on Tests 1-4 (Mean event correlation coefficient of Test 1 amplitude vs mean preference over Tests 1-4, R = 0.320; p = 0.227). **e.** Correlation between event amplitude on Test 1 and preference for the cocaine zone on Test 1 (R = 0.093; p = 0.733). **f.** Correlation between event amplitude on Test 1 and preference for the cocaine zone on Test 2 (R = 0.259; p = 0.333). **g.** Correlation between event amplitude on Test 1 and preference for the cocaine zone on Test 3 (R = 0.392; p = 0.133). **h.** Correlation between event amplitude on Test 1 and preference for the cocaine zone on Test 4 (R = 0.266; p = 0.319).

**Supplementary figure 7.**
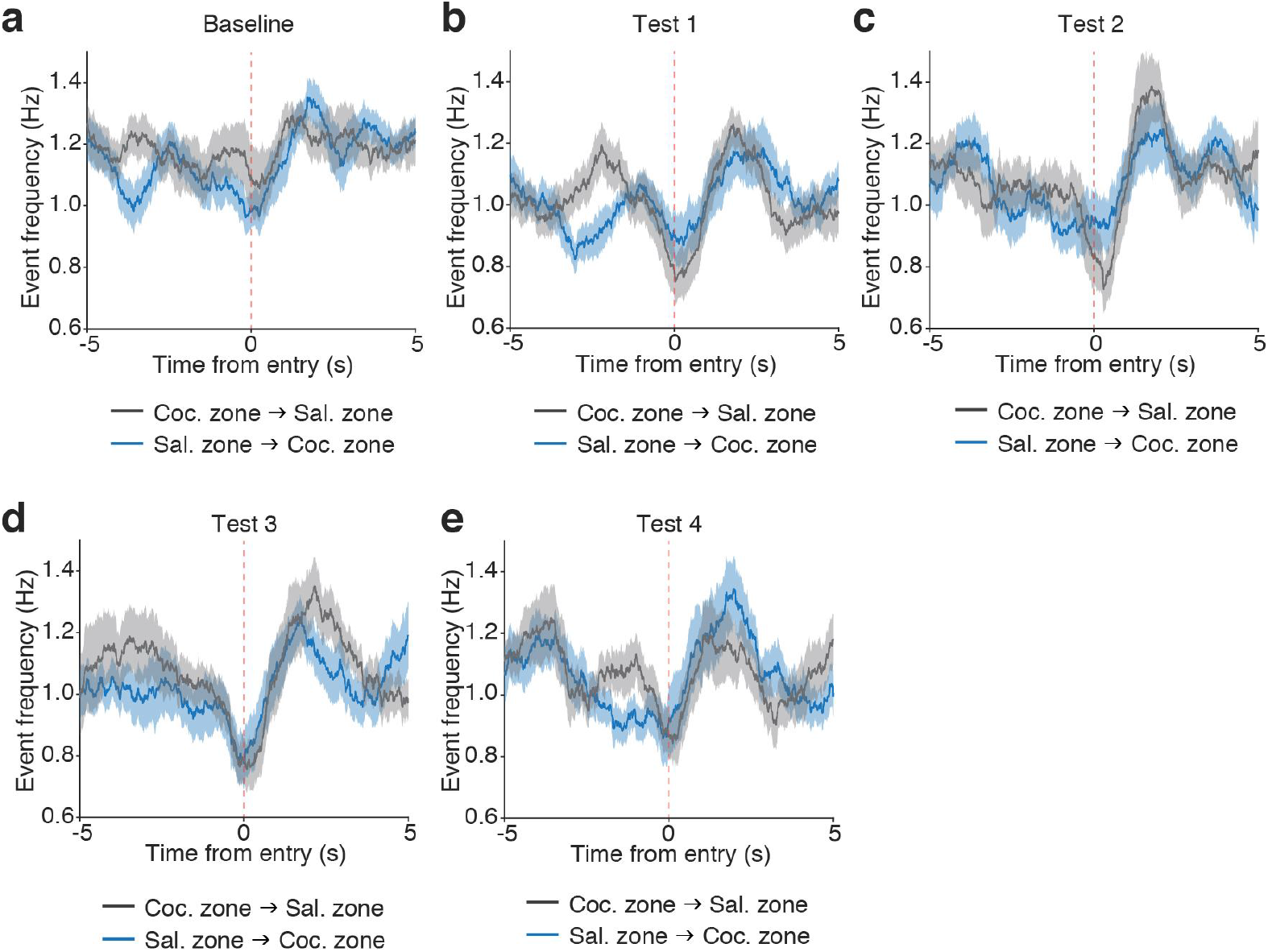
Modulation of ACh event frequency at time of saline and cocaine zone entries across behavioral testing. Mean event frequency around the time of entries into the saline zone (dark grey) and cocaine zone (blue) for Baseline (**a**), Test 1 (**b**), Test 2 (**c**), Test 3 (**d**), and Test 4 (**e**). Event frequency is calculated with a 1 s sliding window over every entry type for each mouse, then averaged across mice.

**Supplementary figure 8.**
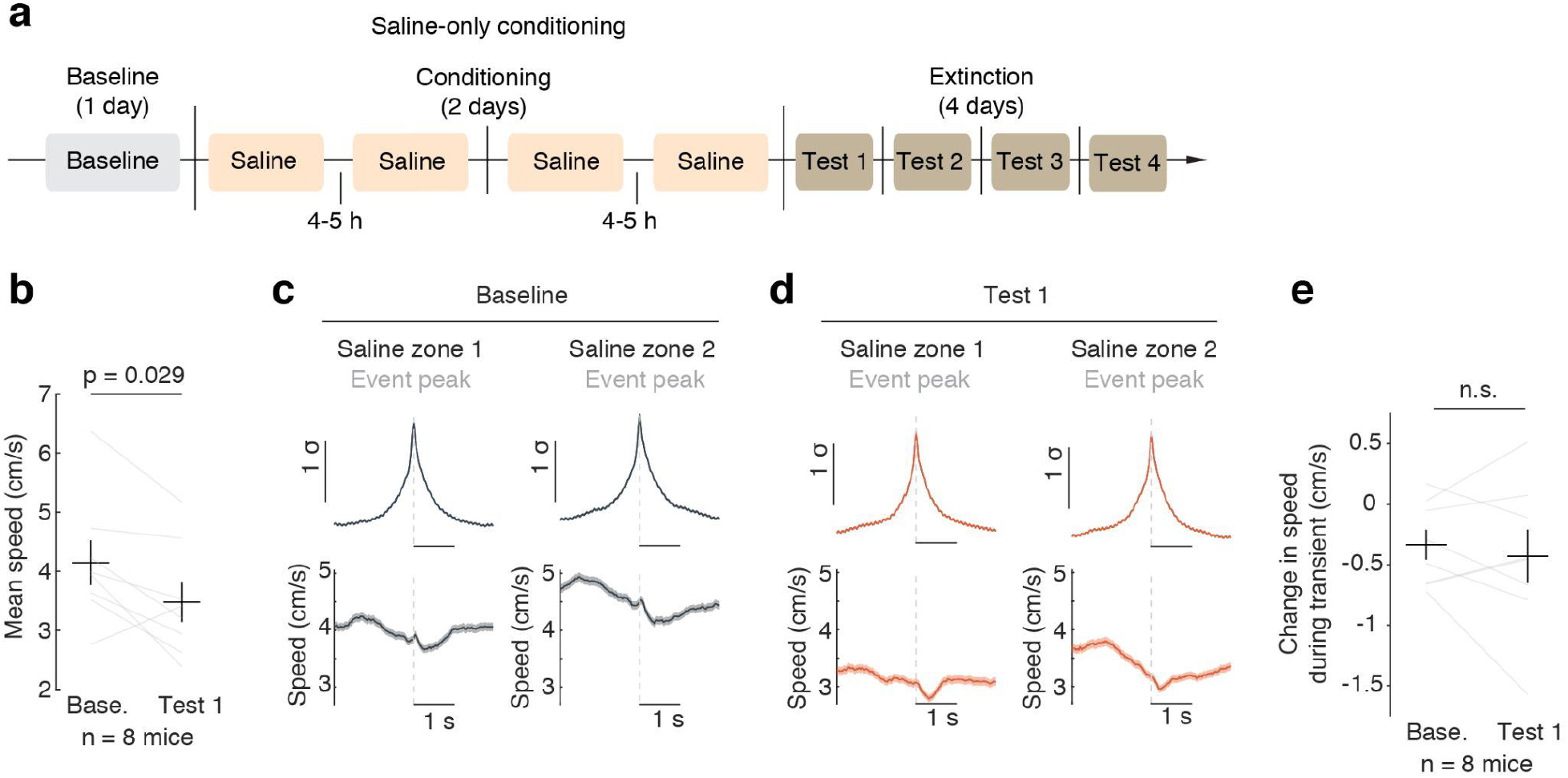
Saline-only conditioning does not affect the relationship between ACh events and speed. **a.** Timeline of saline-only CPP experiment. **b.** Mice show decreased speed on Test 1 compared to Baseline following saline-only conditioning (p = 0.029 for pairwise t-test of Baseline and Test 1). **c.** Mean trace centered at peak for all significant events in the saline 1 zone (top left) and saline 2 zone (top right), with mean speed traces over the same time period (bottom left and right) during Baseline. **d.** Mean trace centered at peak for all significant events in the saline 1 zone (top left) and saline 2 zone (top right), with mean speed traces over the same time period (bottom left and right) during Test 1. **e.** The changes in speed during an ACh event are not different between Baseline and Test 1 (p = 0.547 for pairwise t-test of Baseline and Test 1). Baseline speed was calculated as 0.5 s window starting 2 s before event peak. Speed during peak was calculated as mean speed in a 1.5 s window beginning 0.5 s before event peak and ending 1 s after event peak.

**Supplementary figure 9.**
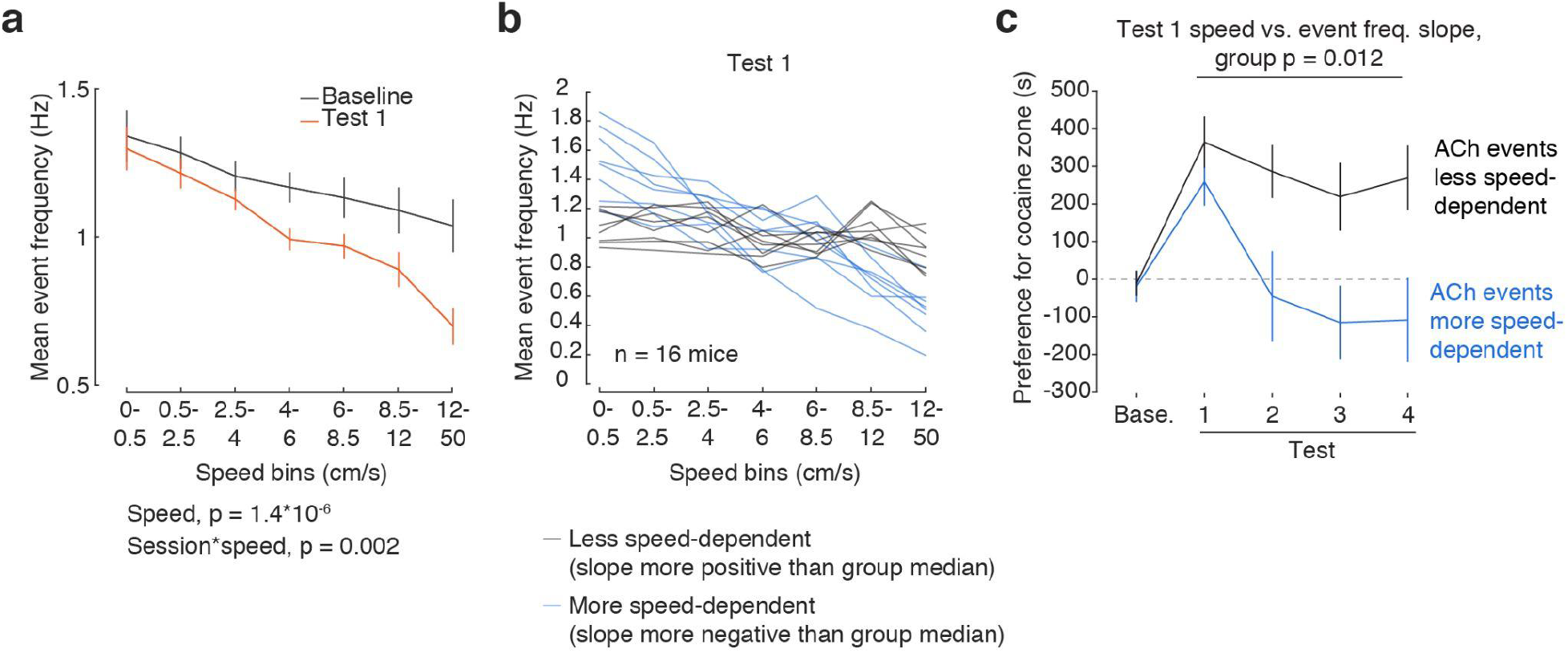
All effects between ACh event frequency, speed, and cocaine zone preference from speed decile-based analyses (Figure 3e-f) are recovered when using defined speed bins. **a.** Mean event frequency by speed during Baseline and Test 1. Event frequency is negatively modulated by speed, and to a greater extent at Test 1 than at Baseline (Effect size d = -0.722, p = 1.4*10^-6^ for speed; effect size d = 0.460, p = 0.002 for session*speed in an LMER with session (Baseline vs Test 1), speed decile, and their interaction as fixed effects and mouse as random effect). **b.** Mean event frequency by speed on Test 1 for individual mice. Individual traces are colored by median split of slope to highlight differences in speed dependency (black: less negative speed vs. frequency slope; blue: more negative speed vs. frequency slope). **c.** Mean preference for the cocaine zone when mice are median split by the slopes of the speed decile vs. Test 1 mean event frequency (Less speed dependence, black, n = 8; More speed dependence, blue, n = 8). Speed decile slope is significantly predictive of preference on Tests 1-4 (F_(1,14)_ = 8.337, p = 0.012 for median split group in repeated measures ANOVA with group, test number, and their interaction as fixed effects and mouse as random effect).

**Supplementary figure 10.**
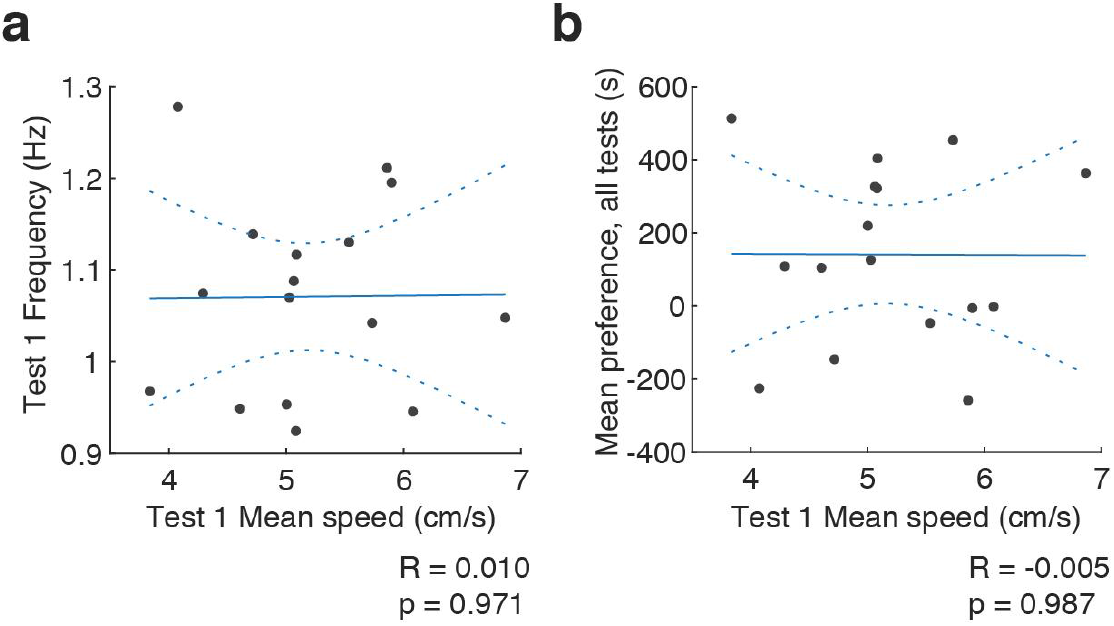
Mean speed on Test 1 does not correlate with mean ACh event frequency on Test 1 or preference for the cocaine zone on Tests 1-4. **a.** Correlation between mean speed on Test 1 and mean ACh event frequency on Test 1 (R = 0.010; p = 0.971). **b.** Correlation between mean speed on Test 1 and mean preference for the cocaine zone across Tests 1-4 (R = -0.005; p = 0.987).

**Supplementary figure 11.**
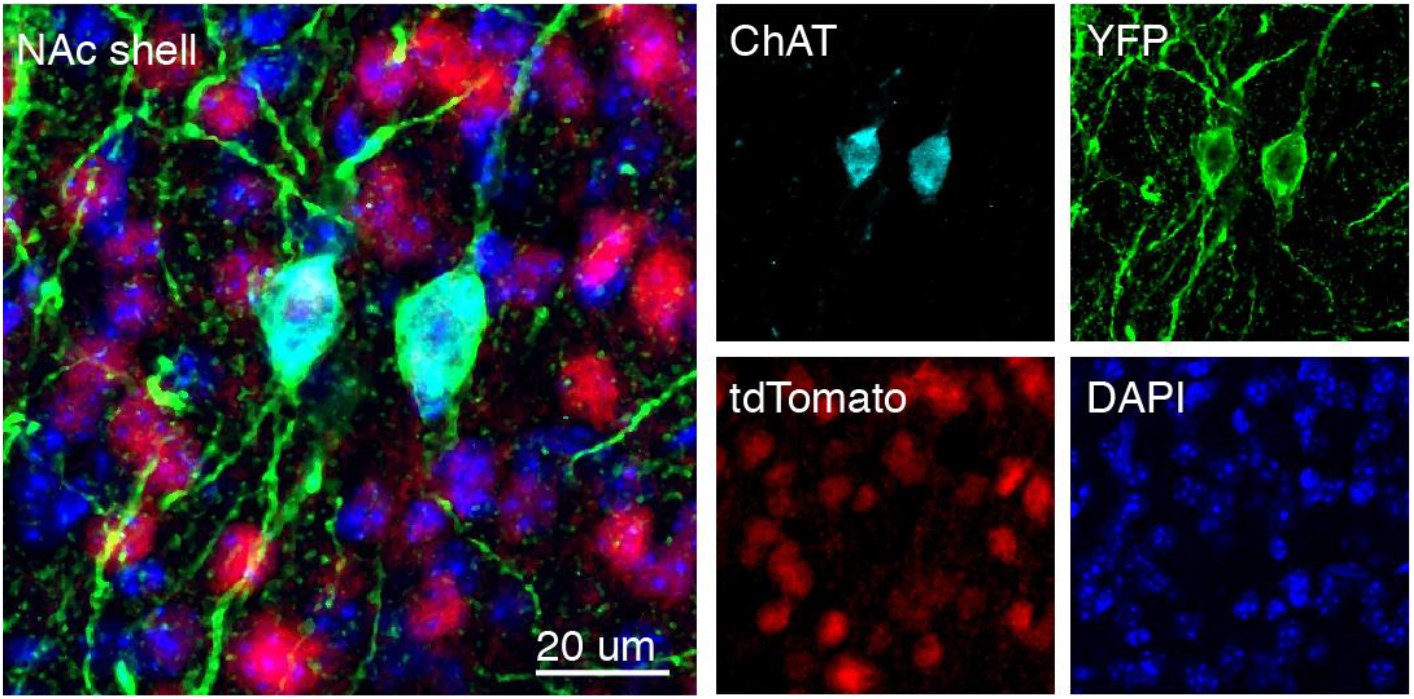
ChR2-YFP expresses selectively in ChINs in ChAT::IRES-Cre / drd1a- tdTomato mice. Co-localization between ChR2-YFP and ChAT immunohistochemistry in NAc shell in tissue from ChAT::IRES-Cre / drd1a-tdTomato mouse. Scale bar, 20 µm.

**Supplementary figure 12.**
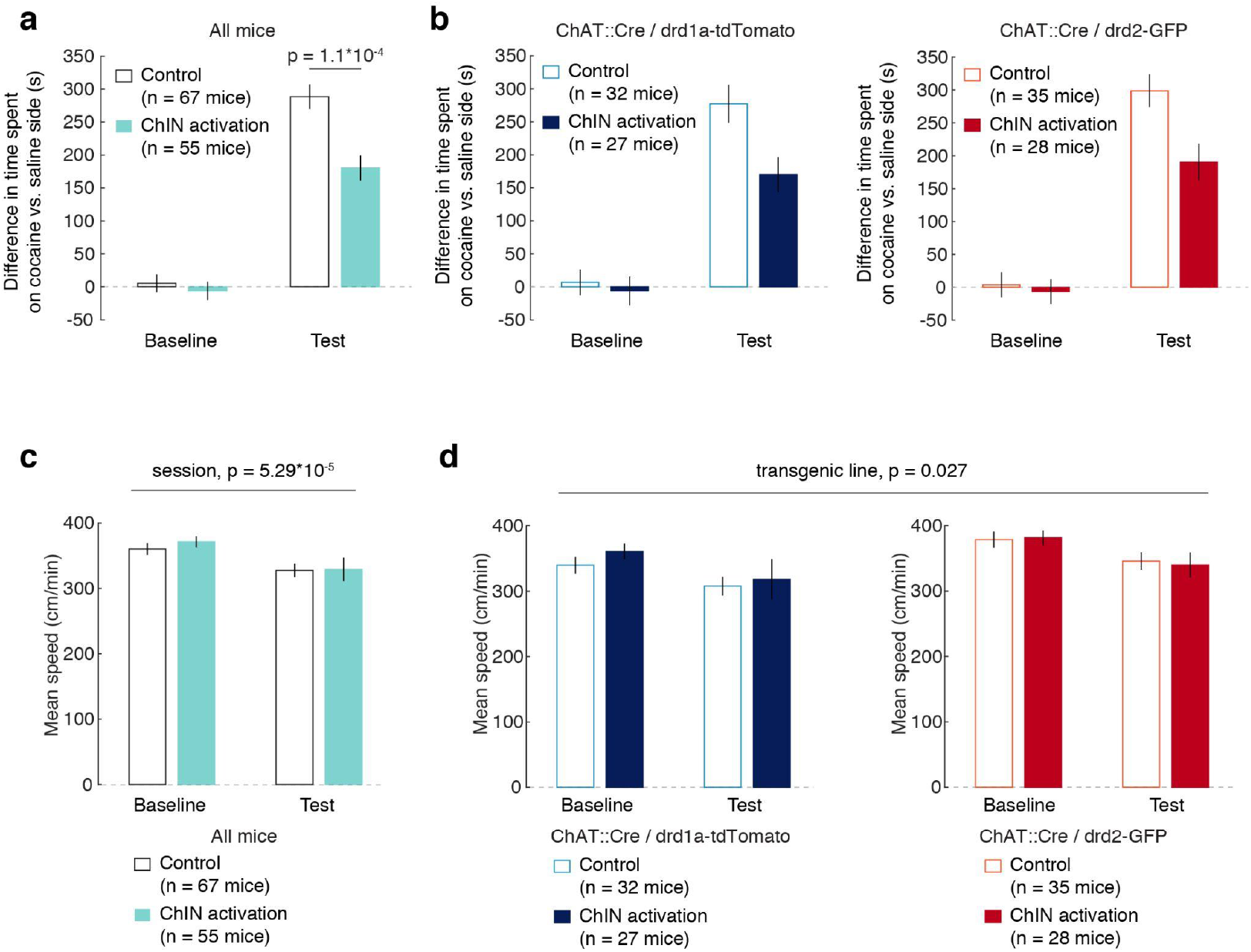
Transgenic line affects meean speed but not CPP formation or ChIN activation. **a-b.** There was no effect of transgenic line on preference, and no interactions between transgenic line and group or between transgenic line and session (F_(1,118)_ = 0.307, p = 0.581 for transgenic line; F_(1,118)_ = 0.689, p = 0.408 for transgenic line*session; F_(1,118)_ = 0.001, p = 0.981 for transgenic line*group; F_(1,118)_ = 0.005, p = 0.942 for transgenic line*session*group; Multi-factor, repeated measures ANOVA with transgenic line (ChAT::Cre/drd1a-tdTomato vs ChAT::Cre/drd2-GFP), group (ChIN activation vs control), and session (Baseline vs Test) as factors.) **c-d.** There were significant effects of session and transgenic line on mean speed, but no effect of ChIN activation and no interactions (F_(1,237)_ = 8.79, p = 0.003 for transgenic line; F_(1,237)_ = 24.7, p < 1*10^-5^ for session; F_(1,237)_ = 0.46, p = 0.499 for group; F_(1,237)_ = 1.75; p = 0.187 for session*group; F_(1,237)_ = 1.53, p = 0.217 for transgenic line*session; F_(1,237)_ = 0.74, p = 0.391 for transgenic line*group; Multi-factor, repeated measures ANOVA with transgenic line (ChAT::Cre/drd1a- tdTomato vs ChAT::Cre/drd2-GFP), group (ChIN activation vs control), session (Baseline vs Test), and their interactions as fixed effects and mouse as random effect.)

**Supplementary figure 13.**
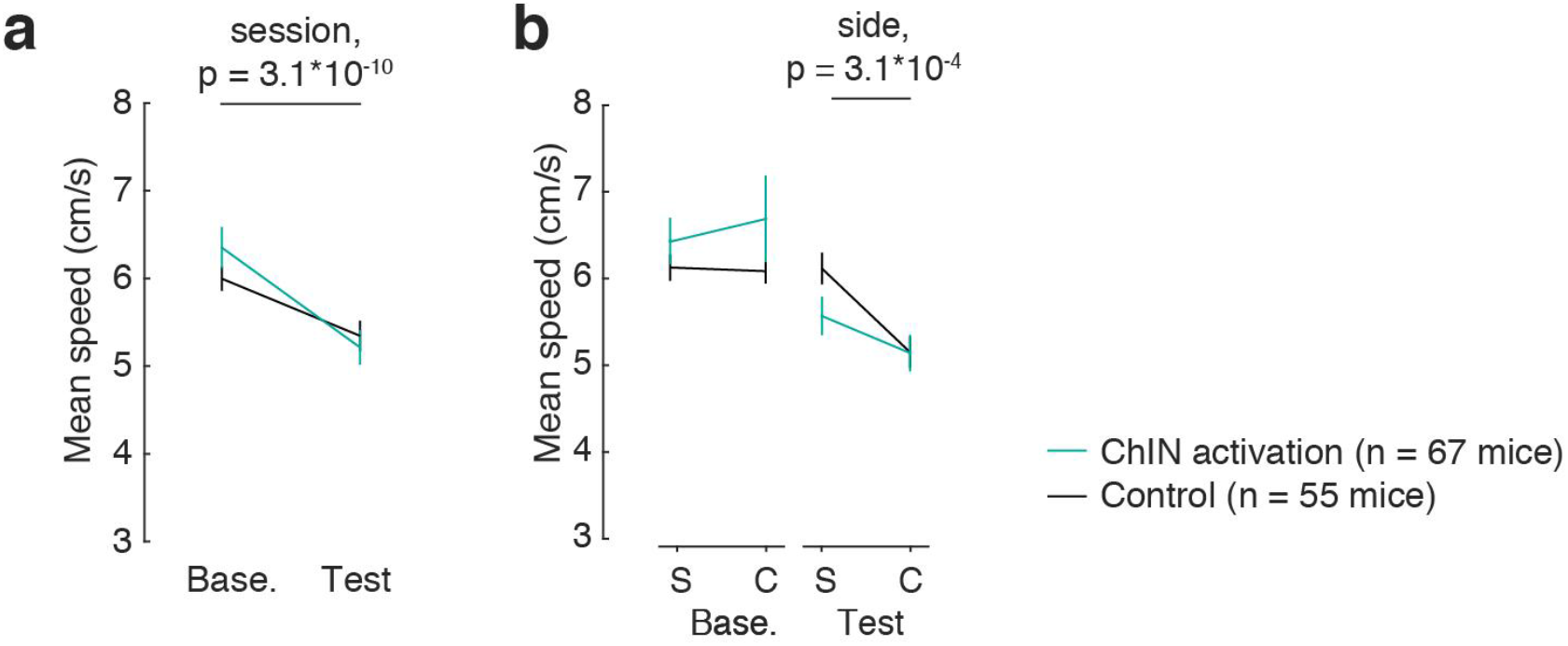
Mice exhibit significantly lower mean speeds on Test in the optogenetic experiment, and ChIN activation does not affect speed by chamber. **a.** Mean speed during the Baseline and Test sessions. Mice are slower on Test compared to Baseline (F_(1,359)_ = 41.969, p = 3.06*10^-10^ for session in a multi-factor, repeated measures ANOVA on speed with session (Baseline vs Test), group (ChIN activation vs control), side (Saline vs Cocaine), and their interactions as fixed effects and mouse as random effect.) **b.** Mean speed by chamber side during Baseline and Test 1. Mice are significantly slower in the cocaine zone compared to the saline zone during the extinction Test (F_(1,239)_ = 13.426, p = 3.1*10^-4^ for side in a multi-factor, repeated measures ANOVA on Test speed with group (ChIN activation vs control), side (Saline vs Cocaine), and their interaction as fixed effect and mouse as random effect), and ChIN activation does not affect speed by chamber side on either session (F_(1,239)_ = 2.653, p = 0.105 for group; F_(1,239)_ = 0.283, p = 0.595 for group*side in a multi-factor, repeated measures ANOVA on Baseline speed with group, side, and their interaction as fixed effect and mouse as random effect; F_(1,239)_ = 1.968, p = 0.162 for group; F_(1,239)_ = 1.854, p = 0.175 for group*side in a multi-factor, repeated measures ANOVA on Test speed with group, side, and their interaction as fixed effect and mouse as random effect).

**Supplementary figure 14.**
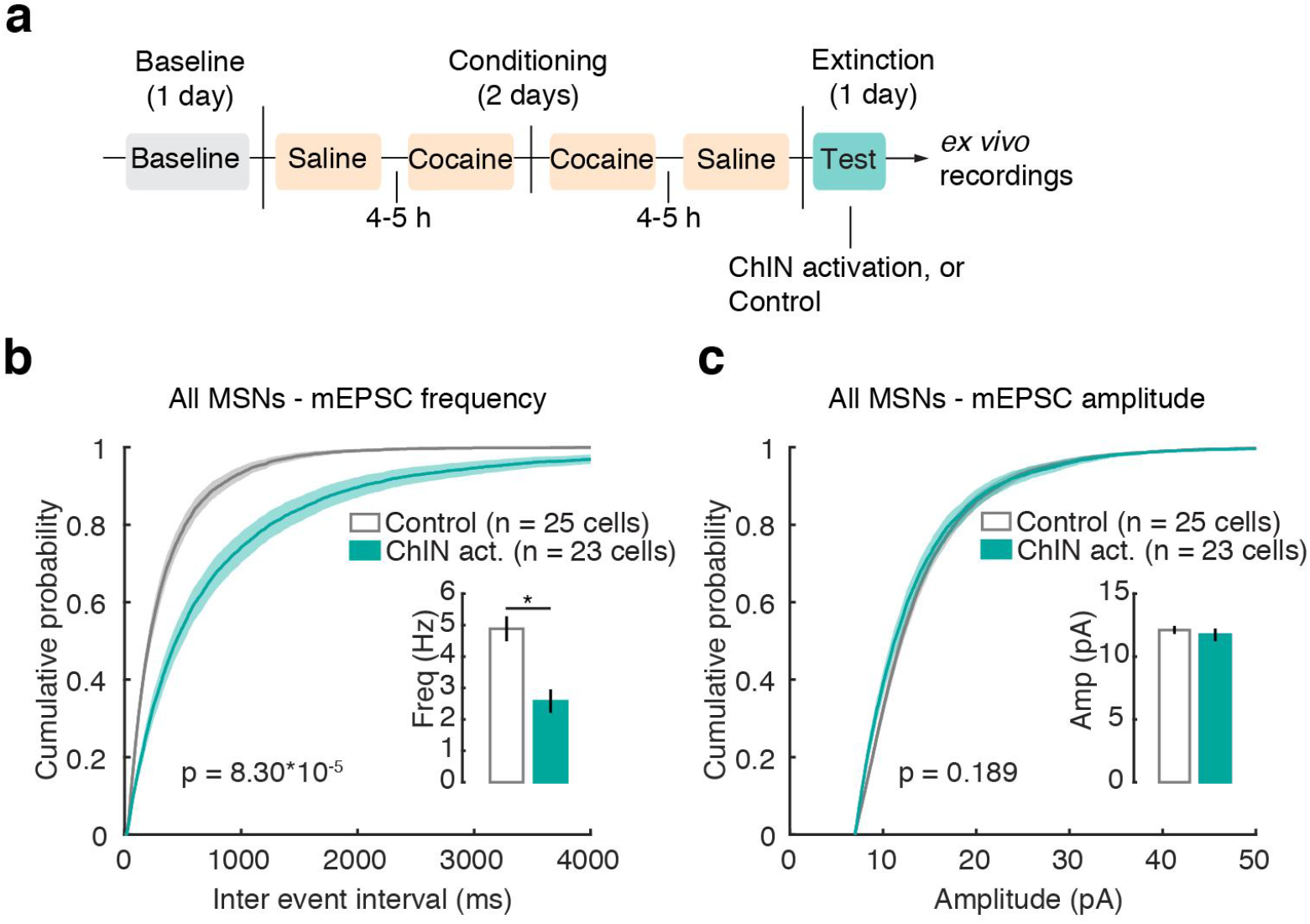
Combined D1R MSN and D2R MSN mEPSC data from Figure 4. Combined data recovers effect of ChIN activation on mEPSC frequency, but masks effect of ChIN activation on mEPSC amplitudes at D2R MSNs. **a.** Timeline of CPP experiments prior to *ex vivo* recordings. One group received optogenetic stimulation of ChINs during extinction testing (447 nm, 15 Hz, 5 ms pulse duration, 2 s light on interleaved with 2 s light off). Immediately after the test, brain slices were collected for *ex vivo* recordings. **b.** Cumulative probability of interevent intervals for mEPSCs at all MSNs. ChIN activation decreased mEPSC frequency at MSNs compared to controls (p = 8.38 * 10^-5^ for group in a LMER on frequency with group (ChIN activation vs control) as fixed effect and cell as random effect. Inset: median frequency of mEPSCs. Control group, n = 25 cells, 22 mice, 4.88 ± 0.40 Hz. ChIN activation group, n = 23 cells, 16 mice, 2.58 ± 0.37 Hz). **c.** Cumulative probability of amplitudes for mEPSCs at all MSNs. ChIN activation did not affect mEPSC amplitude in MSNs compared to controls (p = 0.189 for group in an LMER on amplitude with group (ChIN activation vs control) as fixed effect and cell as random effect. Inset: median amplitude of mEPSCs. Control group, n = 25 cells, 22 mice, 12.11 ± 0.32 pA. ChIN activation group, n = 23 cells, 16 mice, 11.72 ± 0.50 pA).

**Supplementary figure 15.**
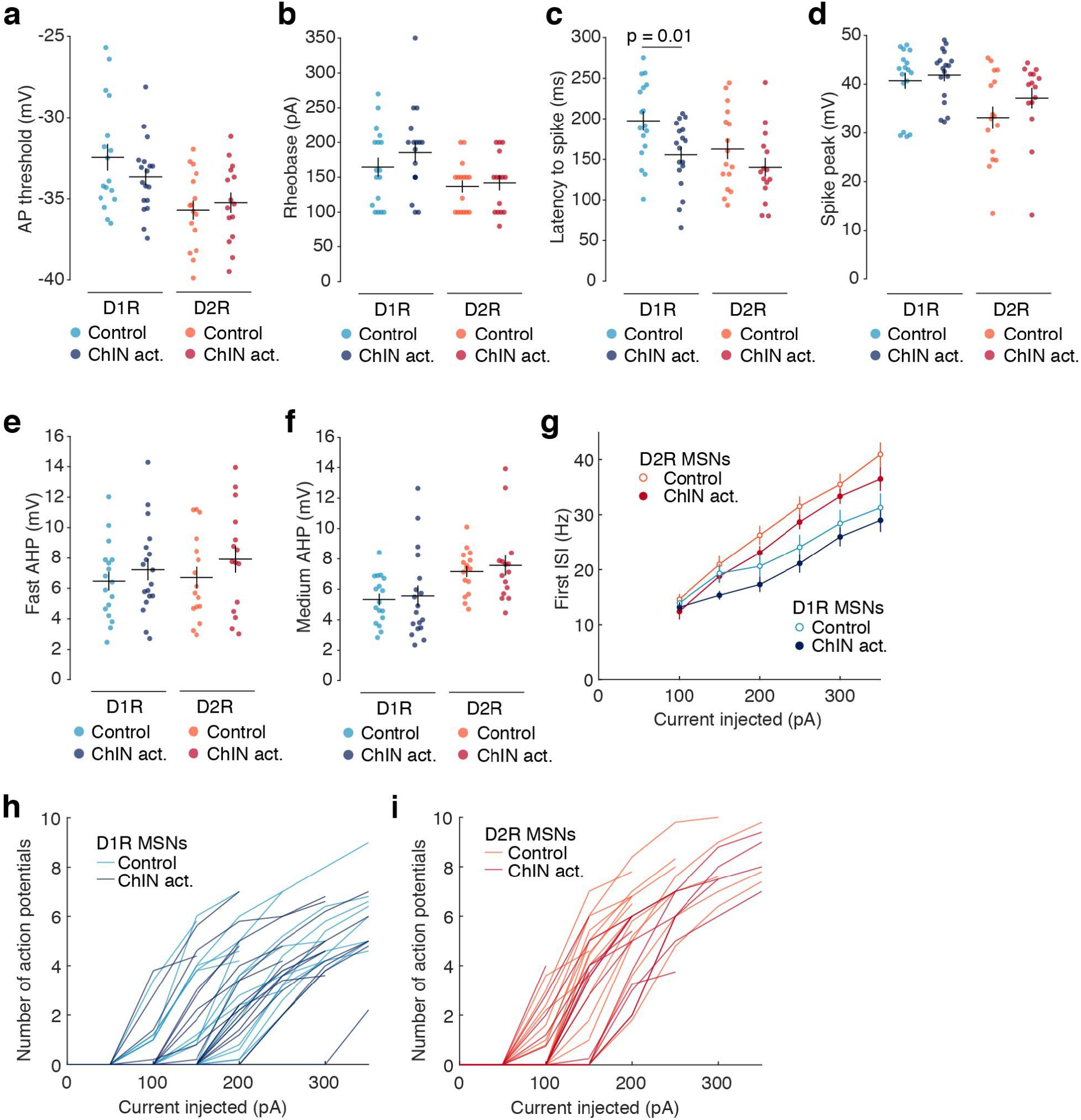
ChIN activation during extinction testing did not affect afterhyperpolarization or spike amplitude statistics. **a.** Summary of mean action potential thresholds. ChIN activation did not significantly alter AP thresholds in D1R MSNs (p = 0.222, two-tailed t test; Control group, n = 17 cells, 5 mice, -32.449 mV ± 0.819. ChIN activation group, n = 18 cells, 4 mice, -33.652 mV ± 0.5336) or D2R MSNs (p = 0.598, two-tailed t test; Control group, n = 17 cells, 7 mice, -35.690 mV ± 0.586. ChIN activation group, n = 15 cells, 4 mice, -35.236 mV ± 0.619). **b.** Summary of mean rheobase currents, measured as the mean of the smallest injected currents to generate a spike on each run of increasing current steps. ChIN activation did not significantly affect rheobase currents in D1R MSNs (p = 0.299, two-tailed t test; Control group, n = 17 cells, 5 mice, 164.706 pA ± 13.480. ChIN activation group, n = 18 cells, 4 mice, 185.556 pA ± 14.376) or D2R MSNs (p = 0.713, two-tailed t test; Control group, n = 17 cells, 7 mice, 136.875 pA ± 8.790. ChIN activation group, n = 15 cells, 4 mice, 142.000 pA ± 10.747). **c.** Summary of mean latency to first spike. ChIN activation significantly decreased latency to spike in D1R MSNs (p = 0.010, two-tailed t test; Control group, n = 17 cells, 5 mice, 197.522 ms ± 11.916. ChIN activation group, n = 18 cells, 4 mice, 155.727 ms ± 9.735), but did not significantly affect latency in D2R MSNs (p = 0.193, two-tailed t test; Control group, n = 17 cells, 7 mice, 162.771 ms ± 12.375. ChIN activation group, n = 15 cells, 4 mice, 140.244 ms ± 11.407). **d.** ChIN activation during extinction testing did not affect spike peak in D1R MSNs (p = 0.581 for two-sample t-test; Control group, n = 16 cells, 5 mice, 40.746 mV ± 1.645. ChIN activation group, n = 18 cells, 4 mice, 41.892 mV ± 1.256) or D2R MSNs (p = 0.209 for two-sample t-test; Control group, n = 16 cells, 7 mice, 33.147 mV ± 2.301. ChIN activation group, n = 15 cells, 4 mice, 37.164 mV ± 2.101). **e.** ChIN activation during extinction testing did not affect fast hyperpolarization in D1R MSNs (p = 0.426 for two-sample t-test; Control group, n = 16 cells, 5 mice, -6.472 mV ± 0.612. ChIN activation group, n = 18 cells, 4 mice, - 7.223 mV ± 0.698) or D2R MSNs (p = 0.294 for two-sample t-test; Control group, n = 16 cells, 7 mice, -6.714 mV ± 0.705. ChIN activation group, n = 15 cells, 4 mice, -7.917 mV ± 0.886). Fast afterhyperpolarization is calculated as the maximum difference between the AP threshold potential and the membrane voltage in an 8 ms window following spike peak. **f.** ChIN activation during extinction testing did not affect medium hyperpolarization in D1R MSNs (p = 0.769 for two-sample t-test; Control group, n = 16 cells, 5 mice, 5.331 mV ± 0.381. ChIN activation group, n = 18 cells, 4 mice, 5.564 mV ± 0.676) or D2R MSNs (p = 0.589 for two-sample t-test; Control group, n = 16 cells, 7 mice, -7.161 mV ± 0.361. ChIN activation group, n = 15 cells, 4 mice, -7.568 mV ± 0.668). Medium afterhyperpolarization is calculated as the maximum difference between the AP threshold potential and the membrane voltage in a 16 ms window following spike peak. **g.** ChIN activation did not affect the first interspike interval (the time between the first two evoked spikes) in D1R MSNs (p = 0.410 for group, p = 0.705 for current *group interaction for D1R MSNs in LMER with group as factor, current as covariate, and cell as random effect) or D2R MSNs (p = 0.703 for group, p = 0.286 for current *group interaction for D2R MSNs in LMER with group as factor, current as covariate, and cell as random effect). i. Number of action potentials evoked as a function of current injected over a 300 ms window in all control (light blue, n = 17 cells) and ChIN activation (dark blue, n = 18 cells) D1R MSNs. Data used to calculate group means in **Figure 4u**. **h.** Number of action potentials evoked as a function of current injected over a 300 ms window in all control (light red, n = 16 cells) and ChIN activation (dark red, n = 15 cells) D2R MSNs. Data used to calculate group means in **Figure 4u**.

## Notes

### Competing Interest Statement

The authors have declared no competing interest.

## References

1. Aitta-Aho, T., Phillips, B.U., Pappa, E., Hay, Y.A., Harnischfeger, F., Heath, C.J., Saksida, L.M., Bussey, T.J., and Apergis-Schoute, J. (2017). Accumbal Cholinergic Interneurons Differentially Influence Motivation Related to Satiety Signaling. eNeuro 4.

2. Al-Hasani, R., Gowrishankar, R., Schmitz, G.P., Pedersen, C.E., Marcus, D.J., Shirley, S.E., Hobbs, T.E., Elerding, A.J., Renaud, S.J., Jing, M., et al. (2021). Ventral tegmental area GABAergic inhibition of cholinergic interneurons in the ventral nucleus accumbens shell promotes reward reinforcement. Nat. Neurosci.

3. Barral, J., Galarraga, E., and Bargas, J. (1999). Muscarinic presynaptic inhibition of neostriatal glutamatergic afferents is mediated by Q-type Ca2+ channels. Brain Res. Bull. 49, 285–289.

4. Baufreton, J., Atherton, J.F., Surmeier, D.J., and Bevan, M.D. (2005). Enhancement of excitatory synaptic integration by GABAergic inhibition in the subthalamic nucleus. J. Neurosci. 25, 8505–8517.

5. Berlanga, M.L., Olsen, C.M., Chen, V., Ikegami, A., Herring, B.E., Duvauchelle, C.L., and Alcantara, A.A. (2003). Cholinergic interneurons of the nucleus accumbens and dorsal striatum are activated by the self-administration of cocaine. Neuroscience 120, 1149–1156.

6. Bernard, V., Normand, E., and Bloch, B. (1992). Phenotypical characterization of the rat striatal neurons expressing muscarinic receptor genes. J. Neurosci. 12, 3591–3600.

7. Bock, R., Shin, J.H., Kaplan, A.R., Dobi, A., Markey, E., Kramer, P.F., Gremel, C.M., Christensen, C.H., Adrover, M.F., and Alvarez, V.A. (2013). Strengthening the accumbal indirect pathway promotes resilience to compulsive cocaine use. Nat. Neurosci. 16, 632–638.

8. Britt, J.P., Benaliouad, F., McDevitt, R.A., Stuber, G.D., Wise, R.A., and Bonci, A. (2012). Synaptic and behavioral profile of multiple glutamatergic inputs to the nucleus accumbens. Neuron 76, 790–803.

9. Cachope, R., Mateo, Y., Mathur, B.N., Irving, J., Wang, H.-L., Morales, M., Lovinger, D.M., and Cheer, J.F. (2012). Selective activation of cholinergic interneurons enhances accumbal phasic dopamine release: setting the tone for reward processing. Cell Rep. 2, 33–41.

10. Cai, L.X., Pizano, K., Gundersen, G.W., Hayes, C.L., Fleming, W.T., Holt, S., Cox, J.M., and Witten, I.B. (2020). Distinct signals in medial and lateral VTA dopamine neurons modulate fear extinction at different times. Elife 9.

11. Calipari, E.S., Bagot, R.C., Purushothaman, I., Davidson, T.J., Yorgason, J.T., Peña, C.J., Walker, D.M., Pirpinias, S.T., Guise, K.G., Ramakrishnan, C., et al. (2016). In vivo imaging identifies temporal signature of D1 and D2 medium spiny neurons in cocaine reward. Proc. Natl. Acad. Sci. U. S. A. 113, 2726–2731.

12. Choi, S., and Lovinger, D.M. (1997). Decreased probability of neurotransmitter release underlies striatal long-term depression and postnatal development of corticostriatal synapses. Proc. Natl. Acad. Sci. U. S. A. 94, 2665–2670.

13. Chuhma, N., Mingote, S., Moore, H., and Rayport, S. (2014). Dopamine neurons control striatal cholinergic neurons via regionally heterogeneous dopamine and glutamate signaling. Neuron 81, 901–912.

14. Cole, S.L., Robinson, M.J.F., and Berridge, K.C. (2018). Optogenetic self-stimulation in the nucleus accumbens: D1 reward versus D2 ambivalence. PLoS One 13, e0207694.

15. Collins, A.L., Aitken, T.J., Huang, I.-W., Shieh, C., Greenfield, V.Y., Monbouquette, H.G., Ostlund, S.B., and Wassum, K.M. (2019). Nucleus Accumbens Cholinergic Interneurons Oppose Cue-Motivated Behavior. Biol. Psychiatry 86, 388–396.

16. Ding, J.B., Guzman, J.N., Peterson, J.D., Goldberg, J.A., and Surmeier, D.J. (2010). Thalamic gating of corticostriatal signaling by cholinergic interneurons. Neuron 67, 294–307.

17. Dorst, M.C., Tokarska, A., Zhou, M., Lee, K., Stagkourakis, S., Broberger, C., Masmanidis, S., and Silberberg, G. (2020). Polysynaptic inhibition between striatal cholinergic interneurons shapes their network activity patterns in a dopamine-dependent manner. Nat. Commun. 11, 5113.

18. Durieux, P.F., Bearzatto, B., Guiducci, S., Buch, T., Waisman, A., Zoli, M., Schiffmann, S.N., and de Kerchove d’Exaerde, A. (2009). D2R striatopallidal neurons inhibit both locomotor and drug reward processes. Nat. Neurosci. 12, 393–395.

19. English, D.F., Ibanez-Sandoval, O., Stark, E., Tecuapetla, F., Buzsáki, G., Deisseroth, K., Tepper, J.M., and Koos, T. (2011). GABAergic circuits mediate the reinforcement-related signals of striatal cholinergic interneurons. Nat. Neurosci. 15, 123–130.

20. Fleming, W., Jewell, S., Engelhard, B., Witten, D.M., and Witten, I.B. (2021). Inferring spikes from calcium imaging in dopamine neurons. PLoS One 16, e0252345.

21. Gallo, E.F., Meszaros, J., Sherman, J.D., Chohan, M.O., Teboul, E., Choi, C.S., Moore, H., Javitch, J.A., and Kellendonk, C. (2018). Accumbens dopamine D2 receptors increase motivation by decreasing inhibitory transmission to the ventral pallidum. Nat. Commun. 9, 1086.

22. Gerfen, C.R., Engber, T.M., Mahan, L.C., Susel, Z., Chase, T.N., Monsma, F.J., Jr, and Sibley, D.R. (1990). D1 and D2 dopamine receptor-regulated gene expression of striatonigral and striatopallidal neurons. Science 250, 1429–1432.

23. Guzman, S.J., Schlögl, A., and Schmidt-Hieber, C. (2014). Stimfit: quantifying electrophysiological data with Python. Front. Neuroinform. 8, 16.

24. Hernández-Echeagaray, E., Galarraga, E., and Bargas, J. (1998). 3-Alpha-chloro-imperialine, a potent blocker of cholinergic presynaptic modulation of glutamatergic afferents in the rat neostriatum. Neuropharmacology 37, 1493–1502.

25. Higley, M.J., Soler-Llavina, G.J., and Sabatini, B.L. (2009). Cholinergic modulation of multivesicular release regulates striatal synaptic potency and integration. Nat. Neurosci. 12, 1121–1128.

26. Hikida, T., Kimura, K., Wada, N., Funabiki, K., and Nakanishi, S. (2010). Distinct roles of synaptic transmission in direct and indirect striatal pathways to reward and aversive behavior. Neuron 66, 896–907.

27. Howe, M., Ridouh, I., Allegra Mascaro, A.L., Larios, A., Azcorra, M., and Dombeck, D.A. (2019). Coordination of rapid cholinergic and dopaminergic signaling in striatum during spontaneous movement. Elife 8.

28. Iino, Y., Sawada, T., Yamaguchi, K., Tajiri, M., Ishii, S., Kasai, H., and Yagishita, S. (2020). Dopamine D2 receptors in discrimination learning and spine enlargement. Nature 579, 555–560.

29. Jing, M., Li, Y., Zeng, J., Huang, P., Skirzewski, M., Kljakic, O., Peng, W., Qian, T., Tan, K., Zou, J., et al. (2020). An optimized acetylcholine sensor for monitoring in vivo cholinergic activity. Nat. Methods 17, 1139–1146.

30. Juarez, B., Morel, C., Ku, S.M., Liu, Y., Zhang, H., Montgomery, S., Gregoire, H., Ribeiro, E., Crumiller, M., Roman-Ortiz, C., et al. (2017). Midbrain circuit regulation of individual alcohol drinking behaviors in mice. Nature Communications 8.

31. Kourrich, S., Rothwell, P.E., Klug, J.R., and Thomas, M.J. (2007). Cocaine experience controls bidirectional synaptic plasticity in the nucleus accumbens. J. Neurosci. 27, 7921–7928.

32. Kravitz, A.V., Tye, L.D., and Kreitzer, A.C. (2012). Distinct roles for direct and indirect pathway striatal neurons in reinforcement. Nat. Neurosci. 15, 816–818.

33. Lee, J., Finkelstein, J., Choi, J.Y., and Witten, I.B. (2016). Linking Cholinergic Interneurons, Synaptic Plasticity, and Behavior during the Extinction of a Cocaine-Context Association. Neuron 90, 1071–1085.

34. Lee, J.H., Ribeiro, E.A., Kim, J., Ko, B., Kronman, H., Jeong, Y.H., Kim, J.K., Janak, P.H., Nestler, E.J., Koo, J.W., et al. (2020). Dopaminergic Regulation of Nucleus Accumbens Cholinergic Interneurons Demarcates Susceptibility to Cocaine Addiction. Biol. Psychiatry 88, 746–757.

35. Lee, S.J., Lodder, B., Chen, Y., Patriarchi, T., Tian, L., and Sabatini, B.L. (2021). Cell-type- specific asynchronous modulation of PKA by dopamine in learning. Nature 590, 451–456.

36. Lim, S.A.O., Kang, U.J., and McGehee, D.S. (2014). Striatal cholinergic interneuron regulation and circuit effects. Front. Synaptic Neurosci. 6, 22.

37. Lobo, M.K., Covington, H.E., 3rd, Chaudhury, D., Friedman, A.K., Sun, H., Damez-Werno, D., Dietz, D.M., Zaman, S., Koo, J.W., Kennedy, P.J., et al. (2010). Cell type-specific loss of BDNF signaling mimics optogenetic control of cocaine reward. Science 330, 385–390.

38. MacAskill, A.F., Cassel, J.M., and Carter, A.G. (2014). Cocaine exposure reorganizes cell type– and input-specific connectivity in the nucleus accumbens. Nature Neuroscience 17, 1198–1207.

39. Malenka, R.C., and Kocsis, J.D. (1988). Presynaptic actions of carbachol and adenosine on corticostriatal synaptic transmission studied in vitro. J. Neurosci. 8, 3750–3756.

40. Mu, P., Moyer, J.T., Ishikawa, M., Zhang, Y., Panksepp, J., Sorg, B.A., Schlüter, O.M., and Dong, Y. (2010). Exposure to cocaine dynamically regulates the intrinsic membrane excitability of nucleus accumbens neurons. J. Neurosci. 30, 3689–3699.

41. Müller, C., and Remy, S. (2013). Fast micro-iontophoresis of glutamate and GABA: a useful tool to investigate synaptic integration. J. Vis. Exp.

42. O’Neal, T.J., Nooney, M.N., Thien, K., and Ferguson, S.M. (2020). Chemogenetic modulation of accumbens direct or indirect pathways bidirectionally alters reinstatement of heroin-seeking in high- but not low-risk rats. Neuropsychopharmacology 45, 1251–1262.

43. Pakhotin, P., and Bracci, E. (2007). Cholinergic interneurons control the excitatory input to the striatum. J. Neurosci. 27, 391–400.

44. Parker, N.F., Cameron, C.M., Taliaferro, J.P., Lee, J., Choi, J.Y., Davidson, T.J., Daw, N.D., and Witten, I.B. (2016). Reward and choice encoding in terminals of midbrain dopamine neurons depends on striatal target. Nat. Neurosci. 19, 845–854.

45. Pascoli, V., Terrier, J., Espallergues, J., Valjent, E., O’Connor, E.C., and Lüscher, C. (2014). Contrasting forms of cocaine-evoked plasticity control components of relapse. Nature 509, 459– 464.

46. Phillipson, O.T., and Griffiths, A.C. (1985). The topographic order of inputs to nucleus accumbens in the rat. Neuroscience 16, 275–296.

47. Robinson, T.E., Gorny, G., Mitton, E., and Kolb, B. (2001). Cocaine self-administration alters the morphology of dendrites and dendritic spines in the nucleus accumbens and neocortex. Synapse 39, 257–266.

48. Shen, W., Tian, X., Day, M., Ulrich, S., Tkatch, T., Nathanson, N.M., and Surmeier, D.J. (2007). Cholinergic modulation of Kir2 channels selectively elevates dendritic excitability in striatopallidal neurons. Nat. Neurosci. 10, 1458–1466.

49. Shen, W., Flajolet, M., Greengard, P., and Surmeier, D.J. (2008). Dichotomous dopaminergic control of striatal synaptic plasticity. Science 321, 848–851.

50. Sjulson, L., Peyrache, A., Cumpelik, A., Cassataro, D., and Buzsáki, G. (2018). Cocaine Place Conditioning Strengthens Location-Specific Hippocampal Coupling to the Nucleus Accumbens. Neuron 98, 926–934.e5.

51. Sugita, S., Uchimura, N., Jiang, Z.G., and North, R.A. (1991). Distinct muscarinic receptors inhibit release of gamma-aminobutyric acid and excitatory amino acids in mammalian brain. Proc. Natl. Acad. Sci. U. S. A. 88, 2608–2611.

52. Tang, K., Low, M.J., Grandy, D.K., and Lovinger, D.M. (2001). Dopamine-dependent synaptic plasticity in striatum during in vivo development. Proc. Natl. Acad. Sci. U. S. A. 98, 1255–1260.

53. Threlfell, S., Lalic, T., Platt, N.J., Jennings, K.A., Deisseroth, K., and Cragg, S.J. (2012). Striatal dopamine release is triggered by synchronized activity in cholinergic interneurons. Neuron 75, 58–64.

54. Willett, J.A., Cao, J., Dorris, D.M., Johnson, A.G., Ginnari, L.A., and Meitzen, J. (2019). Electrophysiological Properties of Medium Spiny Neuron Subtypes in the Caudate-Putamen of Prepubertal Male and Female Drd1a-tdTomato Line 6 BAC Transgenic Mice. Eneuro 6, ENEURO.0016–0019.2019.

55. Willuhn, I., Burgeno, L.M., Groblewski, P.A., and Phillips, P.E.M. (2014). Excessive cocaine use results from decreased phasic dopamine signaling in the striatum. Nat. Neurosci. 17, 704–709.

56. Witten, I.B., Lin, S.-C., Brodsky, M., Prakash, R., Diester, I., Anikeeva, P., Gradinaru, V., Ramakrishnan, C., and Deisseroth, K. (2010). Cholinergic interneurons control local circuit activity and cocaine conditioning. Science 330, 1677–1681.

57. Yan, Z., Flores-Hernandez, J., and Surmeier, D.J. (2001). Coordinated expression of muscarinic receptor messenger RNAs in striatal medium spiny neurons. Neuroscience 103, 1017–1024.

58. Yorgason, J.T., Zeppenfeld, D.M., and Williams, J.T. (2017). Cholinergic Interneurons Underlie Spontaneous Dopamine Release in Nucleus Accumbens. J. Neurosci. 37, 2086–2096.

